# Cryo-EM structures of G4-stalled CMG reveal inchworm mechanism of DNA translocation

**DOI:** 10.1101/2024.10.02.616340

**Authors:** Sahil Batra, Benjamin Allwein, Charanya Kumar, Sujan Devbhandari, Jan-Gert Bruning, Soon Bahng, Chong Lee, Kenneth J. Marians, Richard K. Hite, Dirk Remus

## Abstract

DNA G-quadruplexes (G4s) are non-B-form DNA secondary structures that threaten genome stability by impeding DNA replication. To elucidate how G4s induce replication fork arrest, we have characterized fork collisions with preformed G4s in the parental DNA using fully reconstituted yeast and human replisomes. We demonstrate that a single G4 in the leading strand template is sufficient to stall replisomes by blocking the CMG helicase. An ensemble of high-resolution cryo-EM structures of stalled yeast and human CMG complexes reveals that the G4 is fully folded and lodged inside the CMG central channel. The stalled CMG is conformationally constrained and arrests in the transition between translocation states. Unexpectedly, our analysis suggests that CMG employs an unprecedented inchworm mechanism to translocate on DNA. These findings illuminate the eukaryotic replication fork mechanism under both normal and perturbed conditions.

## Introduction

Faithful replication of the chromosomal DNA prior to cell division is essential for the maintenance of genome integrity. Chromosomal DNA replication in all cells occurs at replication forks and is catalyzed by large, multi-subunit protein complexes, called replisomes, which coordinate the unwinding of the parental DNA with the synthesis of the two daughter DNA strands. Importantly, uninterrupted replisome progression is frequently challenged by obstacles in the chromosomal template, such as transcription, DNA damage, R-loops or non-B-form DNA secondary structures, causing replication stress that poses a threat to genome stability (*1*).

DNA G-quadruplexes (G4s) are particularly abundant non-B-form DNA secondary structures that can form at telomeres and during the transcription or replication of G-rich repeats across the genome (*2*). G4s are polymorphic four-stranded structures comprising a stack of G-quartets that are each composed of four guanines arranged in a planar ring via Hoogsteen hydrogen-bonded base pairing (*3*). While implicated in physiological functions, such as transcriptional regulation, telomere maintenance and immuno-globulin class switching, genomic G4s can also pose an obstacle to DNA replication by impeding DNA polymerases (*4*) or the replicative DNA helicase, CMG (<u>C</u>dc45-<u>M</u>CM-<u>G</u>INS) (*5, 6*). G4s formed during DNA replication are primarily encountered by DNA polymerases in the wake of the CMG (*7, 8*), whereas G4s formed during transcription will be initially encountered by the CMG (*5, 9*). Notably, G4 persistence is not dependent on ongoing transcription (*10, 11*) and R-loops likely play a prominent role in stabilizing G4s as suggested by the strong overlap between G4s and R-loops (*10, 12-15*). Consistent with this notion, G4-stabilizing ligands induce R-loop-dependent transcription-replication conflict in human cells (*16, 17*) and G4s contribute to the block to fork progression at R-loops *in vitro* (*5, 9*). Due to their genotoxic potential, G4 formation is tightly controlled by a large cohort of G4-binding proteins and specialized DNA helicases (*18, 19*). Dysregulation of G4 dynamics upon functional loss of proteins that control G4 formation or stabilization of G4s by small molecule ligands threatens genome integrity, in large part by disrupting chromosomal DNA replication (*16, 20-24*). This has prompted investigations into the cancer therapeutic potential of G4 ligands (*25, 26*).

The CMG is composed of a hetero-hexameric ring of Mcm2-7 (MCM) ATPase subunits and the non-catalytic subunits, Cdc45 and GINS (*27*). The MCM ring forms a two-tiered structure comprising a structural N-terminal tier (N-tier) and a catalytic C-terminal tier (C-tier) harboring the AAA+ ATPase domains. At the replication fork, CMG encircles and translocates in the 3’-5’ direction along the leading strand template while sterically displacing the lagging strand template (*28*). Recent cryogenic electron-microscopy (cryo-EM) studies have resolved structures of CMG bound to forked DNAs, revealing details of the protein-DNA contacts that mediate DNA unwinding and guide the DNA template through the helicase channel (*29-33*). Thus, β-hairpin loops inside the N-tier coordinate the fork junction, whereas ATP-dependent DNA translocation is mediated by the C-terminal pre-sensor 1 (PS1) and helix-2-insert (H2I) β-hairpin loops that protrude from the AAA+ domains into the central channel. Despite these advances, the mechanism of DNA translocation by the CMG, like that of other hexameric ring helicases, remains incompletely understood (*34, 35*). Consequently, competing CMG translocation models have been put forward. In the most commonly accepted rotary hand-over-hand model, ATP turnover proceeds sequentially around the MCM ring with individual MCM subunits concomitantly advancing in 3’-5’ direction along the DNA substrate analogous to a circular staircase (*30, 36*). To accommodate the fact that individual MCM ATPase sites are dispensable for CMG helicase activity (*30, 37*), an asymmetric hand-over-hand mechanism has been proposed in which specific subunits advance pairwise along the DNA (*30*). In contrast to the rotational model, an alternative “pumpjack” model suggested that CMG cycles between extended and compact states during translocation, driven by alternations in N-tier/C-tier distance (*38*). However, since this model was based on CMG structures obtained in the absence of DNA, the relevance of the pumpjack motion for DNA translocation was subsequently questioned (*32*).

In the replisome, CMG associates with a host of accessory proteins, including the fork protection complex (FPC), composed of Mrc1 and Tof1-Csm3 in budding yeast or CLASPIN and TIMELESS(TIM)-TIPIN in humans, which promotes normal fork progression and enhances the stability of stalled forks (*39*). Tof1-Csm3/TIM-TIPIN form a crescent-shaped α-helical complex that associates with the MCM N-tier and coordinates approximately one turn of parental DNA ahead of the CMG (*29, 31, 33, 40*). The extensive engagement of parental DNA by the CMG-FPC both upstream and downstream of the fork junction raises the question where the G4-dependent block to fork progression occurs in the replisome. To address this question, we have examined collisions of fully reconstituted yeast and human replisomes with preformed G4s in the parental DNA and determined an ensemble of high-resolution cryo-EM structures of yeast and human CMGs stalled at a leading strand G4. The data demonstrate a conserved mechanism underlying G4-induced replisome stalling and, unexpectedly, suggest a novel DNA translocation mechanism for CMG.

## Results

### A single G4 in the leading strand template is a potent block to the yeast replisome

In a previous study, using the reconstituted, origin-dependent budding yeast DNA replication system, we observed that R-loop-associated G4s on the leading strand template impede replication fork progression by inhibiting the DNA unwinding activity of CMG (*5*). A limitation of this study was the structural heterogeneity of the R-loop templates, which resulted in variable G4 formation on the displaced strands (*5*). Therefore, to investigate the impact of preformed G4s on fork progression under controlled conditions, we generated ∼7 kbp DNA templates harboring a single, structurally defined G4 positioned ∼1.9 kbp from a yeast replication origin. These templates were generated by ligating DNA fragments containing a preformed G4 site-specifically into plasmid DNA, analogous to a previous approach (*6*). Subsequent linearization of the templates places the G4 in the path of the rightward replication fork while the leftward fork runs off the opposite template end. We focused our studies on a prototypical G4, composed of the sequence (GGGT)_4_, which forms a parallel propeller-type structure (*41*). To stabilize the G4, a poly(dT) sequence was placed opposite the G4.

As expected, a random B-form DNA fragment does not present an impediment to the rightward fork (**Figure 1**). A minor population of leading strand products terminates at the position of the inserted DNA fragment, which is attributed to replisome run-off at leading strand nicks resulting from incomplete ligation of the DNA insert. In striking contrast, the rightward fork was efficiently stalled at a preformed G4 in the leading strand template as evidenced by the accumulation of stalled leading strand products at the expense of full-length leading strand products in the denaturing gel and the concomitant accumulation of stalled fork structures in the native gel. Uncoupled replication products in the native gel indicate limited replisome bypass of the G4 in the absence of continued leading strand synthesis. Consistent with earlier observations (*5, 6*), a preformed G4 on the lagging strand template did not block fork progression. Thus, a single preformed G4 specifically on the leading strand template is sufficient to stall yeast replication forks.

**Figure 1:**
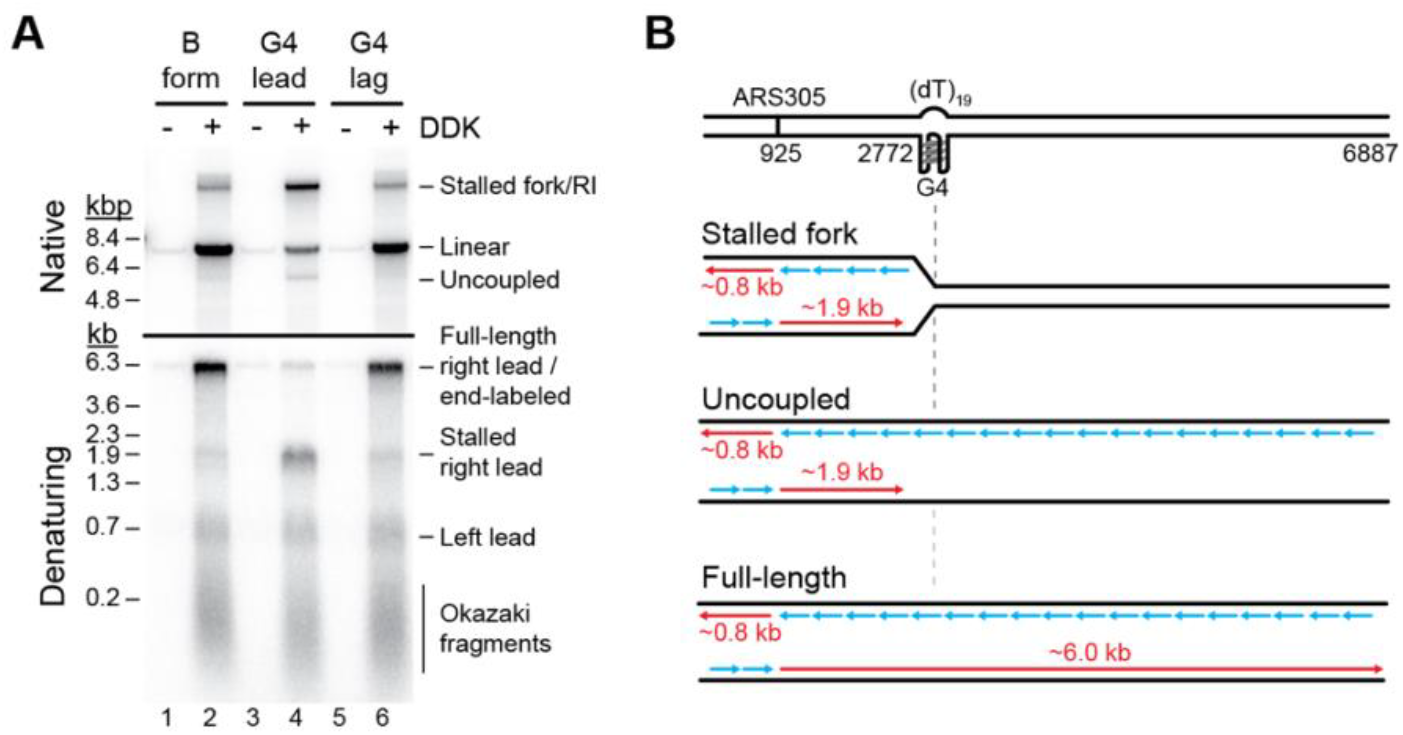
A single G4 in the leading strand template is sufficient to block replication fork progression. **(A)** Denaturing (bottom) and native (top) agarose gel analysis of replication products obtained in the reconstituted yeast system on DNA templates harboring no G4 (lanes 1+2), a G4 on the leading strand template (lanes 3+4) or a G4 on the lagging strand template (lanes 5+6). **(B)** Schematic of replication products obtained in (A). Red: Leading strand products; Blue: Lagging strand products.

Since the FPC has been implicated in the replication of structured DNA repeats (*42*) and G4 sequences (*43*), we asked if the FPC influences CMG helicase activity on G4-containing DNA templates. For this, CMG helicase assays were conducted in the absence or presence of Mrc1/Tof1-Csm3. Consistent with previous observations in the origin-dependent yeast DNA replication system (*44, 45*), we find that the FPC promotes both the rate and amplitude of template unwinding by purified CMG, demonstrating that the stimulatory effect of the FPC on CMG helicase activity is direct (**Figure S1**).

Moreover, like CMG alone (*5, 46*), CMG-FPC is inhibited specifically by a G4 in the leading strand template (**Figure S1**). This indicates that the FPC is neither responsible for the G4-induced block to CMG progression nor does it promote the unwinding of G4-containing DNA by CMG.

To determine if the block to CMG progression is influenced by G4 topology, we tested the effect of G4s with parallel, antiparallel or hybrid topologies (*47*), as well as a G4 derived from the human CEB25 minisatellite sequence that features an extended 9 nt loop (*48*), on CMG-FPC helicase activity. All tested G4s inhibited DNA unwinding by CMG-FPC to a similar degree (**Figure S2**). Thus, the block to CMG-FPC progression is not dependent on G4 topology.

### The stalled CMG-FPC engages DNA on the 3’ and 5’ sides of a leading strand G4

We next developed exonuclease protection assays to investigate the position of the G4 with respect to CMG-FPC using forked oligonucleotide-based DNA substrates harboring a prototypical G4 24 bp downstream of the fork junction in the leading strand template, followed by 35 bp of dsDNA downstream of the G4. Since the CMG travels in 3’-5’ direction on the leading strand template with the MCM N-tier facing forward (*49, 50*), we mapped the position of the C-terminal face of the CMG with Exonuclease T (Exo T), a single strand-specific 3’-5’ exonuclease (*51*). For this, we radiolabled the 5’ end of the G4-containing strand. Reaction products were analyzed by sequencing gel analysis and phosphor imaging.

The single-stranded leading strand template is efficiently digested by Exo T up to one nucleotide from the 3’ end of the preformed G4, demonstrating that the G4 forms a physical obstacle to Exo T (**Figure S3A**). In contrast, when the leading strand template is annealed to the lagging strand template, Exo T digests the leading strand template only up to the fork junction, consistent with the single strand specificity of Exo T.

To test the utility of our exonuclease protection approach to map the position of CMG-FPC on DNA, we initially probed CMG-FPC stably loaded at the fork junction in the presence of the non-hydrolyzable ATP analog adenylyl-imidodiphosphate (AMP-PNP). For this, reactions were probed with Exo T for two minutes prior to DNA extraction.

These conditions yielded a distribution of Exo T stall sites ∼22-24 nt upstream of the fork junction (**Figure S3B**), which agrees with the structural dimensions of CMG-FPC on DNA (*29, 32, 33*). We also observe a CMG-dependent protection of the 3’ end of the leading strand template. Exo T stall sites at the original fork junction are prominent in all reactions due to incomplete template occupancy by CMG-FPC. Time course analyses reveal that CMG-FPC loading at the fork plateaus after 10 minutes (**Figure S3C**) and is stable even when challenged with Exo T for up to 30 minutes (**Figure S3D**).

To determine the position of CMG-FPC stalled at a G4 in the leading strand template, CMG-FPC was assembled at the fork in the presence of a low concentration of AMP-PNP, followed by addition of Exo T and subsequent activation of the CMG-FPC by addition of an excess of ATP (**Figure 2A**). 2 minutes after addition of ATP, the Exo T stall sites 22-24 nt upstream of the fork junction as well as the fully protected strands diminish and new Exo T stall sites appear ∼15 nt upstream of the fork junction, indicating that the CMG-FPC has moved through the fork junction. From 10 minutes onward, Exo T stall sites upstream of the fork junction are absent and a new population of Exo T stall sites, centered ∼13 nt from the G4 3’ end, becomes apparent and remains stable for the duration of the time course. Since Exo T is unable to progress past the fork junction on free DNA, these data indicate that CMG-FPC has unwound the duplex region downstream of the fork junction, allowing Exo T to digest the leading strand template in its wake. Of note, we observe an equivalent CMG-FPC stall position on DNA templates in which the duplex part between the fork junction and the G4 is increased to 34 bp, demonstrating that the CMG-FPC stall position is independent of the distance of the G4 from the original fork junction (**Figure S4A**). We conclude that the C-terminal face of the stalled CMG is ∼13 nt removed from the 3’ end of the G4, suggesting that the G4 is positioned inside the CMG-FPC.

**Figure 2:**
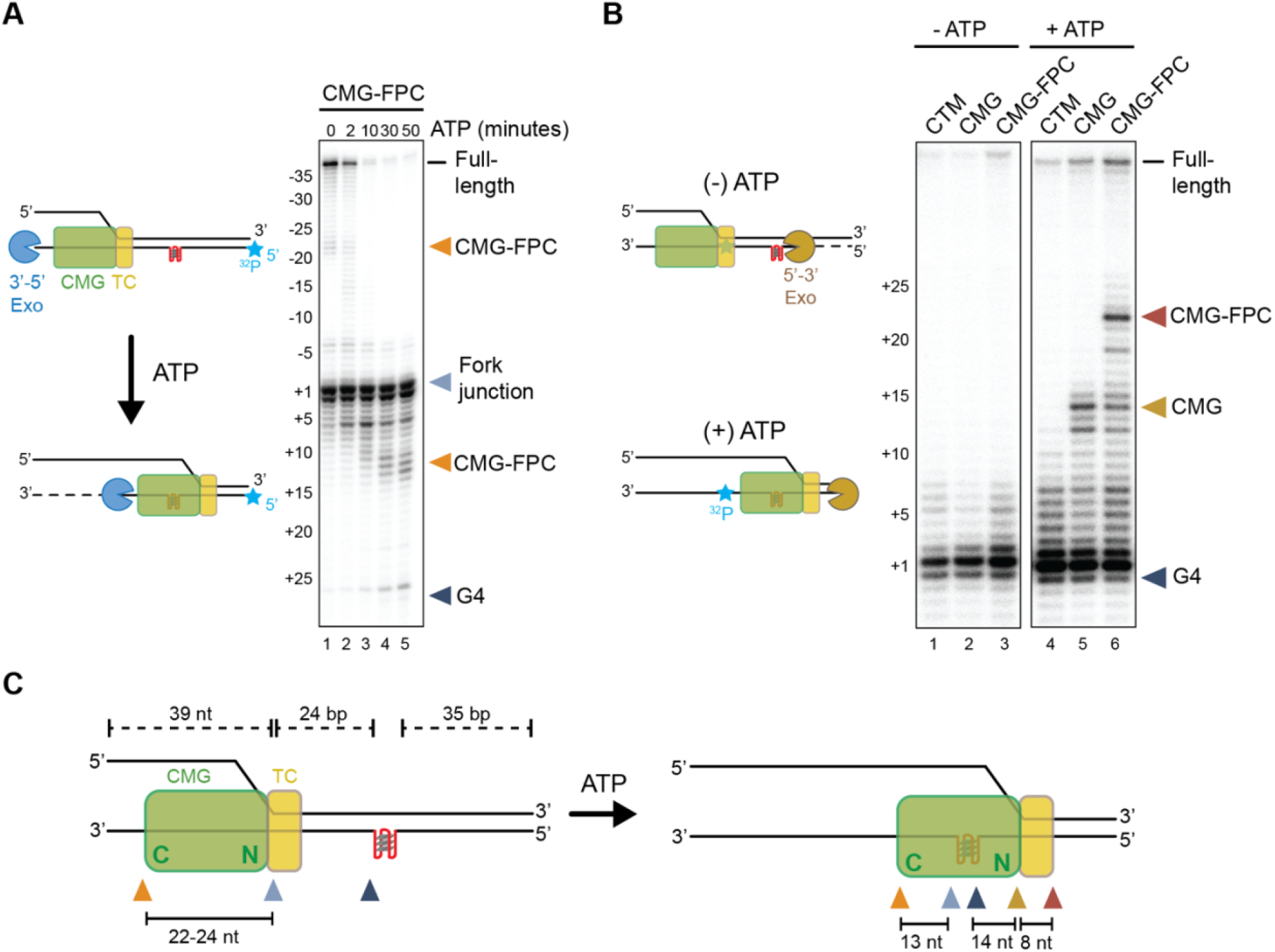
The stalled CMG-FPC protects DNA upstream and downstream of the G4. **(A)** 3’ exonuclease protection analysis to determine position of the C-tier of yeast CMG-FPC stalled at G4 on the leading strand template. **(B)** 5’ exonuclease protection analysis to determine position of the N-tier of yeast CMG-FPC stalled at G4 on the leading strand template. **(C)** Schematic summarizing data from (A) and (B). Green: CMG. Yellow: Tof1-Csm3 (TC). N: N-terminal. C: C-terminal.

If the G4 has entered the CMG channel, DNA downstream of the G4 should be protected by the stalled CMG-FPC. To test this prediction, we developed a 5’ exonuclease protection assay to map the position of the N-terminal face of the CMG-FPC. For this, we incorporated a single ^32^P group via a splint ligation approach into the backbone of the leading strand template 2 nt downstream of the fork junction and probed the position of CMG-FPC with T5 exonuclease or a combination of 5’ exonucleases with complementary substrate specificities, including T7 exonuclease, RecJf, ExoVIII and T5 exonuclease. Both approaches yielded comparable results.

In the presence of AMP-PNP, i.e. when the CMG-FPC complex is stably bound to the fork junction, the leading strand template is digested up to the G4 (**Figure 2B**). Following activation of CMG-FPC with ATP, DNA downstream of the G4 is protected from 5’ exonuclease digestion, indicative of CMG-FPC stalling at the G4. CMG alone protects up to ∼14 bp downstream of the G4 5’ end, while an additional 8-12 bp are protected in the presence of the FPC, consistent with Tof1-Csm3 engaging ∼1 turn of parental duplex DNA ahead of the CMG (*29*). Collectively, the data demonstrate that the CMG-FPC complex protects up to 13 bp of DNA upstream of the G4 3’ end and 22-26 bp DNA downstream of the G4 5’ end when stalled at a leading strand G4, indicating that the G4 is positioned in the CMG central channel during stalling (**Figure 2C**).

### The G4 is flexibly trapped at the N-tier/C-tier interface inside the central CMG channel

To determine the structural basis for the G4-induced block to replisome progression, we collected cryo-EM images of yeast CMG-FPC stalled at a leading strand G4. As before, CMG-FPC complexes were assembled in the presence of AMP-PNP on forked DNA templates harboring a preformed G4 in the leading strand template, 24 bp downstream of the fork junction. DNA unwinding by CMG-FPC was induced by addition of excess ATP. After an incubation time of 10-30 minutes, reactions were vitrified on cryo-EM grids for image collection without addition of any chemical crosslinking agents. Image analysis revealed multiple distinct populations among the particles. Notable differences among the classes relate to the position of the leading strand template in the MCM C-tier, the position of the G4, the ATP-bound state, and the occupancy of Csm3-Tof1 adjacent to the MCM N-tier. Due to their completeness, we will focus our discussion on two major conformations in which the MCM C-tier and Tof1-Csm3 are well resolved, referred to hereafter as states 1 and 2. Reconstruction of state 1 achieved a global resolution of 3.2 Å that was subsequently improved by focused refinements to resolutions of 3.1-3.2 Å whereas state 2 achieved a global resolution of 3.5 Å that was improved to 3.3-4.3 Å via focused refinements (**Figure S5**).

States 1 and 2 share numerous similar features including possessing the forked DNA template, intact MCM rings, CDC45-GINS and Tof1-Csm3 **(Figure 3A-D**). In both states, we also observed densities that allowed unambiguous modeling of the G4 within the central chamber at the interface between the N- and C-tiers of the MCM ring (**Figure 3B,D**). Within the central chamber, the G4s adopt parallel-stranded propeller-type conformations that superimpose well with the NMR structure of an analogous G4 free in solution (**Figure S6**, RMSD ∼1.5 Å). However, the conformations of the DNA within the central chamber differ for the two states. In state 1, the G4 extends towards Mcm2 and Mcm6. In state 2, the G4 is rotated nearly 180° in plane, resulting in it extending towards Mcm3 and Mcm7. The dynamics of the G4 are afforded by the cavity at the N-tier/C-tier interface, which is large enough to accommodate a fully folded G4 in both states (**Figure 3E+F**). Consistent with the notion that the N-tier can accommodate excess DNA, using the 3’ exonuclease protection assay, we find that the CMG-FPC arrests at identical positions at the compact prototypical G4 and the larger CEB25 G4, which harbors an extended 9 bp loop (**Figure S4B**).

**Figure 3:**
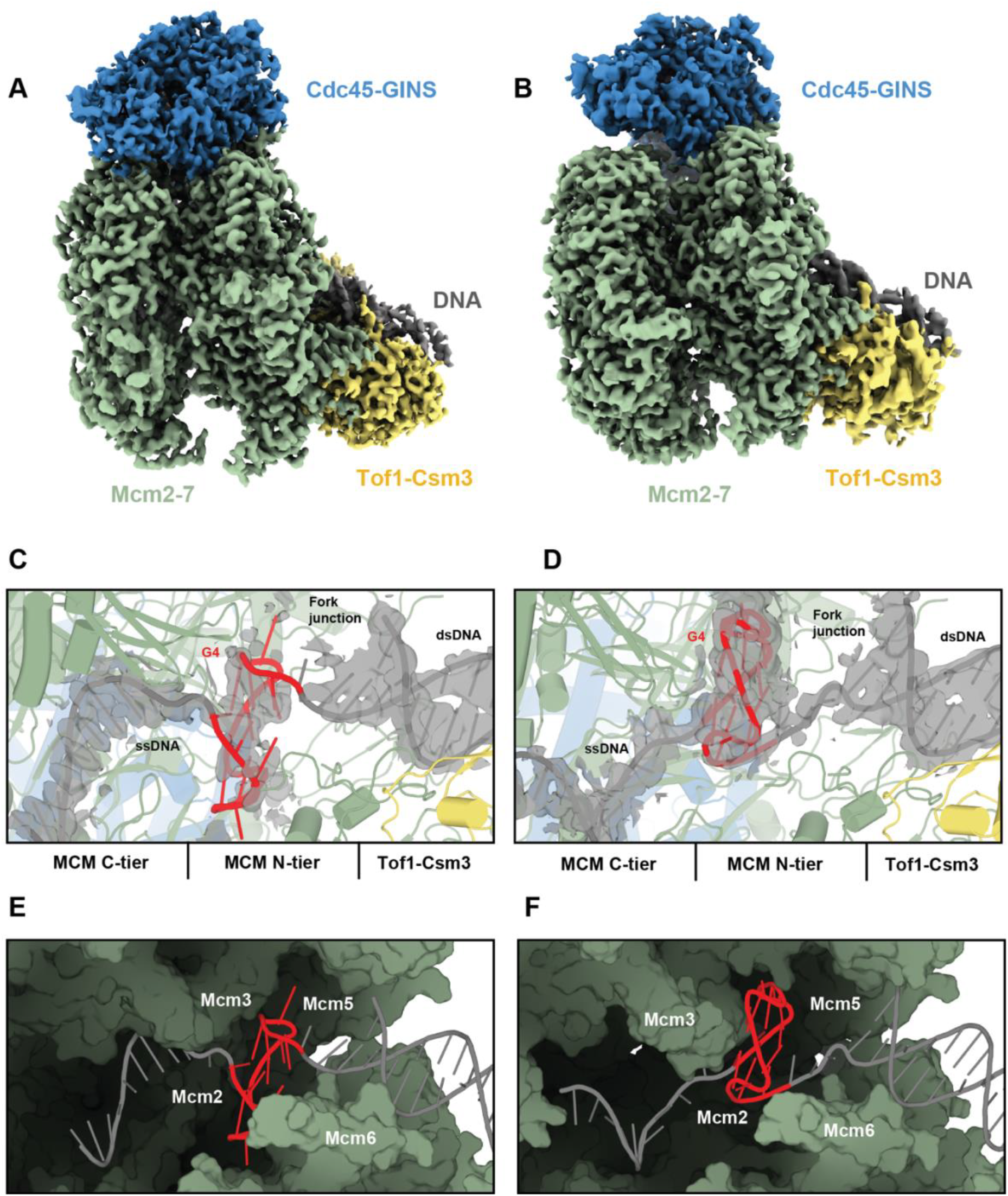
The G4 is lodged inside the central channel of stalled yeast CMG-FPC. **(A,B)** Cryo-EM density map of yeast CMG-FPC stalled at G4, state 1 (A) and state 2 (B) colored by subcomplex with Mcm2-7 in green, Cdc45-GINS in blue and Tof1-Csm3 in yellow. **(C,D)** Structure of the G4 bound in the central chamber of the MCM ring in state 1 (C) and state 2 (D). Densities corresponding to the DNA are shown as grey isosurfaces and contoured at 2.5 s. **(E,F)** Surface representation of MCM central chamber in state 1 (E) and state 2 (F) with Mcm4 and Mcm7 removed for clarity.

Upstream of the G4, the leading strand templates in states 1 and 2 are threaded in a spiral configuration through the C-tier, guided by distinct interactions with the PS1 and H2I hairpin loops emanating from the C-terminal MCM AAA+ domains (**Figure S7**). In state 1, the leading strand template is coordinated in the 5’ to 3’ direction by Mcm3, -5, -2 and -6, whereas Mcm4 and -7 do not engage the DNA. In state 2, the leading strand template associates in 5’ to 3’ direction with Mcm2, -6, -4 and -7, whereas Mcm3 and -5 are unbound from DNA. In both states, the phosphate backbone of the ssDNA is primarily coordinated by highly conserved lysine and alanine residues in the PS1 loop and serine and valine/isoleucine residues at the base of the H2I loop (**Figure S7**). Previous structures of yeast, fly, and human CMG bound to replication forks exhibit nearly identical DNA-protein contacts, indicating that the G4-stalled structures correspond to canonical translocation states (*29-33, 52*). For example, the ssDNA in a structure of yeast CMG-FPC bound to a normal forked DNA (PDB 6SKL), which we will refer to as normal fork conformation 1, is coordinated in a similar manner as the ssDNA in G4 stall state 1. Similarly, the ssDNA in a second conformation bound to a normal fork (PDB 6SKO), which we will refer to as normal fork conformation 2, is coordinated in a similar manner to G4 stall state 2.

Whereas the G4 and the upstream ssDNA adopt distinct conformations in G4 stall states 1 and 2, the conformations of the fork and the dsDNA downstream of the fork are nearly indistinguishable (**Figure 3A-D; Figure S8A**). In both states, the lagging strand template is coordinated by the N-terminal OB-fold hairpin loops of Mcm6, -4 and -7 at the fork junction **(Figure S8B,C)**. These interactions guide the final base pair to the separation pin residue, Mcm7 F363, which enables the diversion of the unwound lagging strand template towards the Mcm3 and Mcm5 Zn^2+^-finger domains. Further downstream of the fork, ∼1 turn of dsDNA is coordinated by the Tof1-Csm3 complex, which tilts the dsDNA in both states at a ∼50º angle relative to the central MCM channel. Notably, the conformations of the fork and downstream dsDNA in states 1 and 2, as well as those of the residues that coordinate them, closely mimic those observed in normal fork conformation 1 **(Figure S8D)** (*29*) and corroborate the 5’ exonuclease protection data **(Figure 2B)**. The striking similarity of the fork observed in the presence of normal forked DNA and forked DNA with a G4 indicates that the G4 does not perturb the protein-DNA contacts that are necessary for DNA unwinding.

### The G4 in the leading strand template impedes the transition between CMG translocation states

At the C-terminal side of the MCM central chamber, the PS1 and H2I loops of Mcm2-7 create a constriction in the central channel. While this constriction accommodates the leading strand template during normal CMG translocation, it is too narrow to accommodate the folded G4 and impedes passage from the N-tier to the C-tier (**Figure 4A+B**). Steric clashes occur between the G4 and the tips of the H2I and PS1 loops (**Figure 4C+D**). In state 1, the G4 contacts the H2I loops of Mcm5 and Mcm2 and the in-plane rotation of the G4 in state 2 orients the G4 towards the H2I loops of Mcm5 and Mcm3.

**Figure 4:**
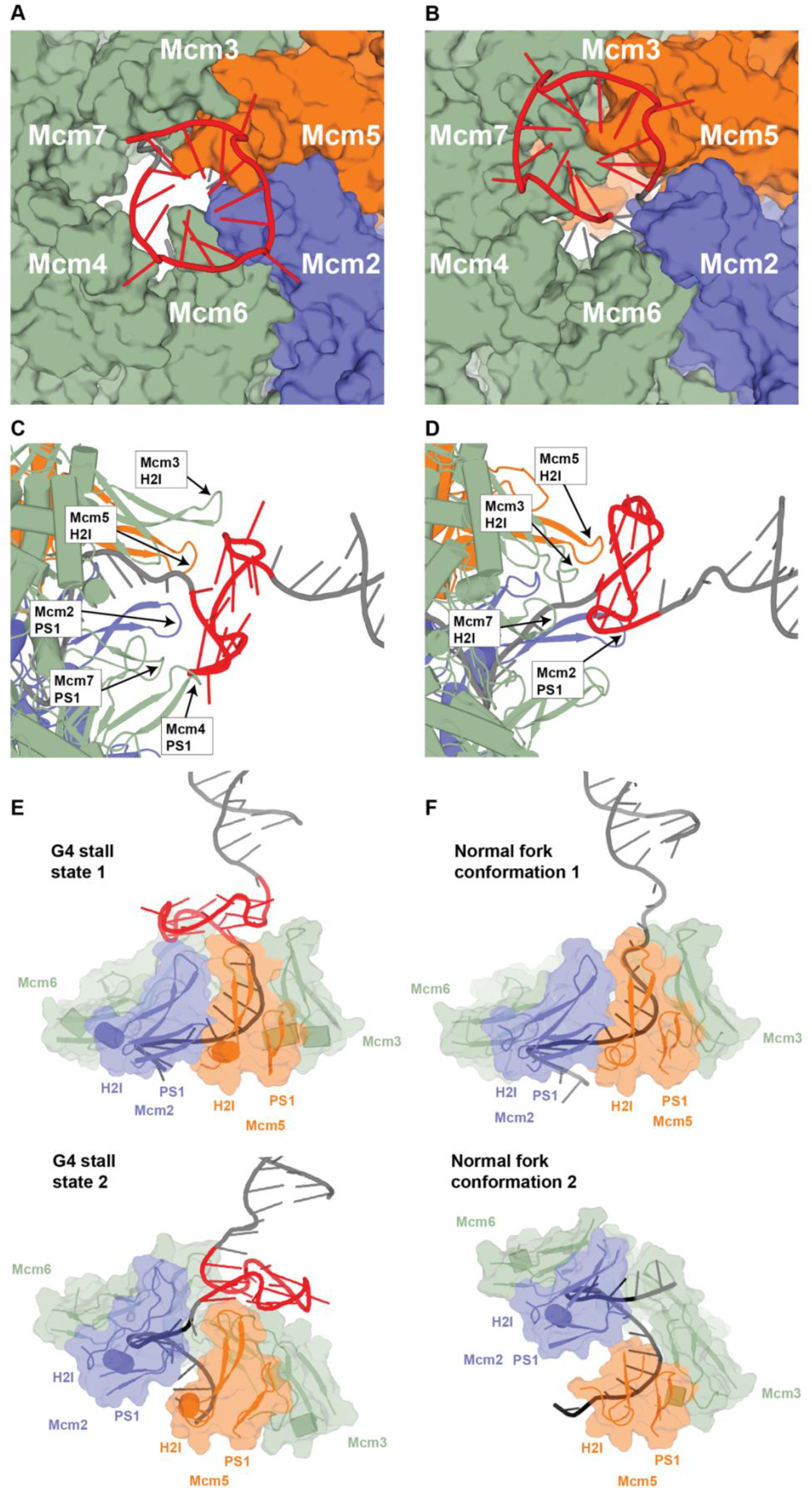
The MCM PS1/H2I loops sterically block passage of the leading strand template at the G4. **(A,B)** Surface representation of the access to the MCM C-tier for state 1 (A) and state 2 (B) with DNA shown as cartoon, viewed from the N-tier. **(C)** Interactions between MCM PS1/H2I loops and the G4 in state 1 (C) and state 2 (D), viewed from the side. **(E-F)** MCM PS1/H2I loops in G4 stall states 1 and 2 (E) and normal fork conformations 1 and 2 (F). Mcm2 is shown in blue, Mcm6, -4, -3, and -7 are shown in green and Mcm5 is shown in orange.

Despite the different orientations of the G4 and the upstream ssDNA in the G4 stall states, the PS1/H2I loops are arranged in a planar manner in both states (**Figure 4E**). In contrast, inspection of structures of yeast and human CMG-FPC bound to normal forked DNAs revealed that translocation of the ssDNA in the MCM C-tier is associated with a rearrangement of the PS1/H2I loops into a spiral configuration (**Figures 4F + S9A**). For example, when the ssDNA is coordinated in normal fork conformation 1 by Mcm3, -5, -2, and -6 in the 5’ to 3’ direction, the PS1/H2I loops adopt a planar ring that is nearly identical to that observed in G4 stall state 1 (**Figure 4E+F**). In normal fork conformation 2, by contrast, the PS1/H2I loops transition to a spiral configuration as DNA is translocated and coordinated by Mcm2, -6, -4, - 7, -3 and -5. Aligning the PS1/H2I loops of Mcm5 in normal fork conformations 1 and 2 reveals that the PS1/H2I loops of Mcm2, -6, -4, -7 and -3, reorient in conformation 2 toward the 5’ proximal side of the leading strand template (**Figure 4F**). The largest movement involves the PS1/H2I domain of Mcm2, which transits ∼20 Å between 3’- and 5’-proximal positions along the channel axis, corresponding to a 12 nt translocation of the leading strand template. The movement of the PS1/H2I loops of Mcm2 in normal forked conformation 2 results in a disruption of the interface between the PS1 loop of Mcm2 and H2 of Mcm5 that is defined by main chain interactions between the highly conserved residues S632/K633 in Mcm2 and G443/S445/G448 in Mcm5 in conformation 1. Equivalent interactions order the remaining MCM interfaces (**Figure S9B,C**).

The transition from the planar to the spiral state clashes with the G4 in state 1, which opposes the Mcm2 PS1/H2I loops. The G4 in stall state 2 also establishes an impediment to the spiral-to-planar transition, resulting in an intermediate state of the translocation process. In this intermediate state, Mcm5 and -3 are released from the DNA at the 3’ proximal end of the spiral and have moved towards the 5’ proximal side of the leading strand template, whereas Mcm2, -6, -4, and partially -7, remain bound to the DNA. However, due to the steric clash between the PS1/H2I loops of Mcm5 and -3 and the G4, the Mcm5 PS1/H2I loops fail to fully engage the PS1/H2I loops of Mcm2 and reestablish 5’ proximal DNA contacts, preventing restoration of the normal planar state and stalling in a quasi-planar state instead (**Figure 4E, S9D,E**).

Our assignment of state 2 as an intermediate in the translocation process implies that the CMG translocates on DNA by alternating between spiral and planar configurations of its DNA-binding PS1/H2I loops, analogous to an inchworm mechanism. In the planar state, represented by G4 stall state 1 and normal fork conformation 1, the Mcm2 and - 5 PS1/H2I loops are physically engaged via conserved mainchain interactions, resulting in a closed-ring conformation.

Transition into the spiral state, which corresponds to normal fork conformation 2, involves the breakage of the Mcm2/5 PS1/H2I loop interface and the movement of the Mcm2 PS1/H2I loops to reengage with the leading strand template 12 nt downstream in the 5’ direction. The interface between the Mcm6 and -2 PS1/H2I loops remains stable during this transition (**Figure S9B,C**), resulting in the concomitant movement of the Mcm6 and -2 PS1/H2I loops. The 3’-5’ movement of the Mcm2 and -6 PS1/H2I loops propagates across the Mcm4, -7 and -3 PS1/H2I loops, allowing the Mcm4 and -7 PS1/H2I loops to engage DNA and culminating in a spiral configuration stabilized by conserved PS1-H2 interactions between all MCM subunits except Mcm2 and -5. Completion of the translocation cycle would involve a sequence of rearrangements, including a transient state analogous to that in G4 stall state 2, that induce the release of the Mcm5 and -3 PS1/H2I loops from the DNA on the 3’ end of the spiral, re-establishment of the PS1-H2 interface between Mcm2 and -5 at the 5’ proximal end of the spiral, re-engagement of the 5’ proximal DNA by the Mcm5 and -3 PS1/H2I loops and release of the Mcm7 and -4 PS1/H2I loops from DNA, thus re-establishing the planar state.

The proposed inchworm mechanism is driven by alternating 12 nt movements of the Mcm2 and -5 PS1/H2I loops and thereby diverges from the previously proposed rotary hand-over-hand mechanism, which postulates a sequential 3’ to 5’ movement of individual MCM subunits or pairs of MCM subunits in 2-4 nt steps around the MCM ring analogous to a circular staircase while maintaining a constitutive planar MCM ring conformation and a largely stable Mcm2/5 interface (*30*). As discussed below, the inchworm mechanism aligns with the asymmetric distribution of ATPase sites essential for CMG helicase activity around the MCM ring, which is more difficult to explain by rotary mechanisms (*30, 37*).

### DNA binding is uncoupled from ATP-binding in the C-tier of the stalled CMG-FPC

To gain further insights into the consequences of CMG-FPC stalling at G4s, we next investigated the ATP binding sites at the interfaces between the six MCM subunits. In G4 stall state 1, ADP is bound at Mcm2; the remaining ATP-binding sites are occupied by ATP. The arrangement of nucleotides is shifted in G4 stall state 2 with ADP being bound at Mcm4 and -6 and ATP being bound at Mcm2, -3, -5 and -7 (**Figure 5A,B**). Notably, the active site configuration at all six AAA+ interfaces in both states 1 and 2 principally corresponds to a catalytically competent state as illustrated by the ordered arrangement of catalytic residues from the Walker A, Walker B and R-finger motifs (**Figure 5C,D**).

**Figure 5:**
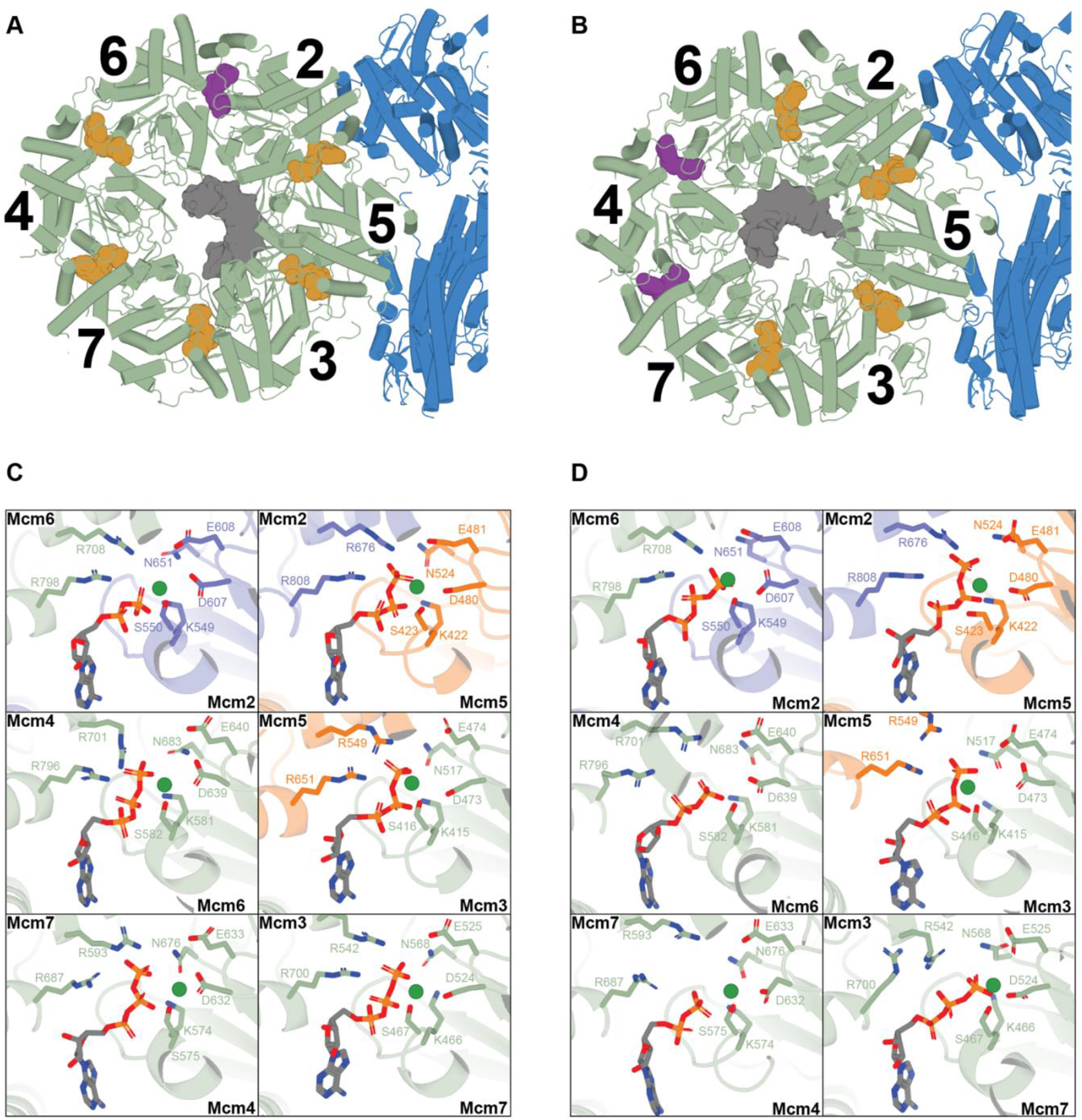
Uncoupling of ATP- and DNA-binding in G4-stalled yeast CMG-FPC. **(A,B)** MCM C-tier of state 1 (A) and state 2 (B), viewed from the N-tier. Bound ATP molecules are shown as orange spheres, bound ADP molecules are shown as magenta spheres and ssDNAs are shown as grey surfaces. **(C,D)** MCM C-tier ATPase active sites in state 1 (C) and state 2 (D).

Although previous studies had indicated that the position of the ssDNA in the C-tier is defined by the ATP-bound state of the AAA+ domains (*29-33, 52*), DNA binding in the C-tier of the G4 stalled CMG-FPC does not strictly correlate with the ATP-bound state of AAA+ domains (**Figure S7**). For example, Mcm4 and -7 in state 1 and Mcm5 and -3 in state 2 are bound to ATP, but do not engage the leading strand template, indicating that conformational constraints associated with CMG arrest can override the DNA binding competence of ATP-bound AAA+ domains. Conversely, Mcm2, -6 and -4 engage the leading strand template in states 1 and 2 in either ADP-or ATP-bound form. While PS1/H2I positions are normally determined by ATP-dependent conformational changes in the C-tier, the PS1/H2I loops of Mcm2:ADP in G4 stall state 1 superimpose on those of Mcm2:AMP-PNP in normal fork conformation 1, while the PS1/H2I loops of Mcm6:ADP and Mcm4:ADP in G4 stall state 2 superimpose on those of Mcm6:AMP-PNP and Mcm4:AMP-PNP in normal fork conformation 2. This indicates that perturbation of the C-tier structure in the stalled CMG-FPC enables DNA binding independent of nucleotide-bound state, i.e. ATP- and DNA-binding are uncoupled.

To determine if the PS1/H2I conformational states described above are associated with global changes in the C-tier, we aligned the MCM C-tiers of CMG-FPC in G4 stall states 1 and 2 on the Mcm5 AAA+ domain. In line with the co-planar arrangement of the PS1/H2I domains in both G4 stall states, the C-tier remains relatively rigid between states 1 and 2 (**Figure S10A**). This contrasts the rearrangement of the AAA+ domains in CMG-FPC bound to normal forked DNA, which are arranged in a co-planar manner in conformation 1 but adopt a spiral configuration in conformation 2, with Mcm2 and -5 marking the endpoints of the spiral (**Figure S10B**). The Mcm2 AAA+ domain undergoes the largest change, shifting by ∼ 17 Å, while Mcm6, -4, -7 and -3 incur progressively smaller shifts, mirroring the co-axial movement of the corresponding PS1/H2I domains. Previous studies also observed rigid body movements of the C-tier, involving the tilting and rotation of the C-tier relative to the N-tier (*29, 33, 38*). In contrast, alignment of G4 stall states 1 and 2 on the N-tier demonstrates that the C-tier is significantly less dynamic in CMG stalled at the G4 (**Figure S10C**).

### The molecular basis for G4-induced fork arrest is conserved in the human replisome

We next investigated if the mechanism of replisome block by G4s is broadly conserved by characterizing the G4-induced fork block using a reconstituted human DNA replication system analogous to that described recently (*53*).

Specifically, replisomes reconstituted from purified human CMG (hCMG), CLASPIN, TIM-TIPIN, AND-1, Pol α, Pol δ, Pol ε, RPA, RFC, PCNA and CTF18-RFC (**Figure S11A**) were assembled in the presence of a low concentration of AMP-PNP on forked DNA templates harboring a preformed G4 1.9 kbp downstream of the fork junction, followed by the induction of fork progression via addition of excess ATP. Analogous to our observations in the yeast system, we find that a single G4 in the leading strand template induces efficient fork stalling, while a G4 in the lagging strand template does not measurably affect fork progression (**Figure 6A,B**). As before, a background of stalled leading strand products is observed on templates harboring a DNA insert due to the retention of nicks in the DNA during template preparation. These background products are not observed on a linear control template prepared without insertion of a DNA fragment by ligation (“un-scarred”). Thus, a single G4 specifically in the leading strand template is a potent block to both yeast and human replisomes.

**Figure 6:**
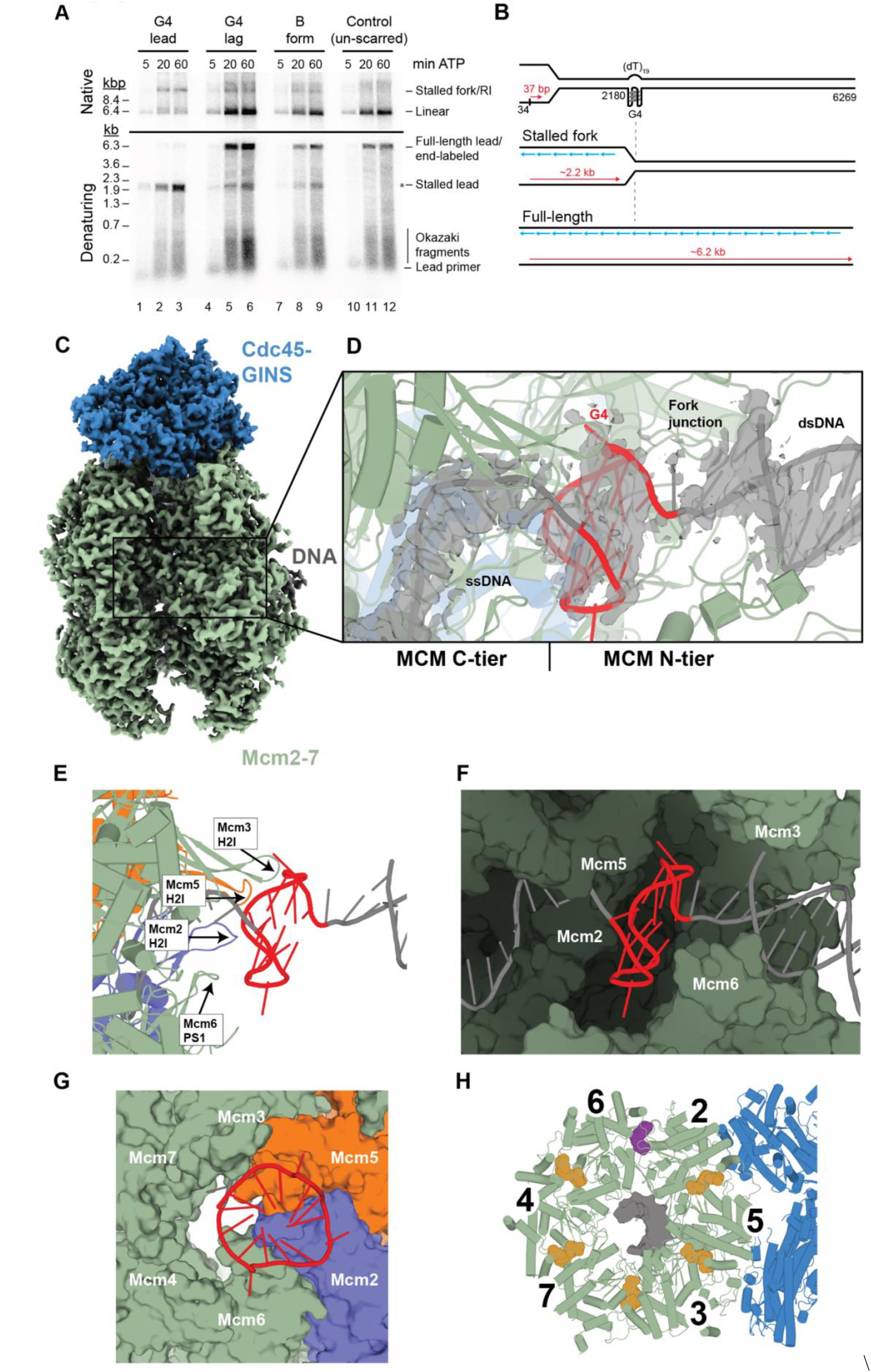
The G4 stall mechanism is conserved in the human replisome. **(A)** Denaturing (bottom) and native (top) agarose gel analysis of replication products obtained in the reconstituted human system on DNA templates harboring no G4 (lanes 1+2), a G4 on the leading strand template (lanes 3+4) or a G4 on the lagging strand template (lanes 5+6). **(B)** Schematic of replication products obtained in (A). **(C)** Cryo-EM density map of human CMG-FPC stalled at G4 colored by subcomplex. **(D)** Structure of the G4 bound in the central chamber of the MCM ring. Density corresponding to the DNA is shown as a grey isosurface and contoured at 2.5 s. **(E)**. Interactions between MCM PS1/H2I loops and bound G4, viewed from the side. **(F)** Surface representation of MCM central chamber with Mcm4 and Mcm7 removed for clarity **(G)** Surface representation of the access to the MCM C-tier with DNA shown as cartoon, viewed from the N-tier. **(H)** MCM C-tier, viewed from the N-tier. Bound ATP molecules are shown as orange spheres, bound ADP molecules are shown as magenta spheres and ssDNAs are shown as grey surfaces.

To determine the structural basis for the G4-induced block to human replisome progression, we characterized hCMG-FPC stalled at a leading strand G4 by cryo-EM. Human CMG-FPC complexes were assembled in the presence of AMP-PNP on forked DNA templates harboring a preformed prototypical G4 in the leading strand template, 24 bp downstream of the fork junction. DNA unwinding was induced by addition of excess ATP. After an incubation time of 10-30 minutes, reactions were vitrified without the addition of chemical crosslinking agents. Image analysis yielded a single conformational state at a global resolution of 2.6 Å that was improved to 2.4 - 2.6 Å using focused refinements **(Figure 6C)**. The DNA is well-defined and can be unambiguously modeled, revealing that the G4 has entered the central channel of the hCMG and is blocked from entering the C-tier at the N-tier/C-tier interface **(Figure 6D-G)**. Remarkably, nearly all of the features resolved in yeast G4 stall state 1 are recapitulated in the DNA in the G4-stalled hCMG, including the conformation of the G4 in the central cavity, the coordination of the upstream ssDNA and the coordination of the downstream fork junction (**Figure S11B,C)**. Both the arrangement of the PS1/H2I domains **(Figure S11D**) and ATPase site occupancy **(Figure 6H)** of yeast G4 stall state 1 are also recapitulated in hCMG. Notably, no density for TIM-TIPIN is observed, suggesting a weaker association of the FPC with hCMG than with yCMG. However, we note that the TIM-TIPIN complex is detected in at least a fraction of the stalled hCMG complexes using the 5’ exonuclease protection assay (**Figure S11E**), indicating that FPC binding to the hCMG is destabilized by cryo-EM conditions.

Collectively, the data demonstrate that the mechanism of G4-induced fork stalling is identical in the yeast and human systems. In both cases, the G4 passes the FPC at the front of the replisome and the fork junction inside the N-tier but is blocked from entering the C-tier by steric clashes with PS1/H2I pore loops, arresting the CMG with the fully folded G4 lodged inside the central channel.

## Discussion

Here, we demonstrate that even a single G4 can create a strong steric block inside the CMG central channel. This provides physical evidence for the replisome blocking potential of G4s, which is often assessed in the presence of G4-stabilizing ligands. Moreover, while G4 stability may be modulated by G4-binding proteins in cellular and extract-based approaches, our utilization of fully reconstituted systems demonstrates that the impact of single G4s on fork progression is intrinsic to the G4 structure. Given the prevalence of potential G-quadruplex sequences (PQSs) across eukaryotic genomes (*54, 55*), this demonstrates that G4s must be continuously turned over during S phase to allow unhindered fork progression, emphasizing the importance of specialized cellular mechanisms to limit G4 formation and maintain genome stability (*19*).

Our data demonstrate that the G4-induced block to replisome progression occurs specifically at preformed G4s in the leading strand template, whereas G4s in the lagging strand template do not impede fork progression. This is consistent with the observation that steric blocks on the lagging strand are generally bypassed by the CMG (*28, 56*). The internalization of the G4 by the CMG, which sequesters the G4 from G4-controlling proteins, has implications for potential fork restart mechanisms at sites of G4-induced fork arrest. For example, the canonical configuration of the fork junction is predicted to protect the replisome from ubiquitin-mediated disassembly (*57*), which may rule out replisome disassembly as a mechanism to expose the G4 for processing by G4 helicases and allow completion of DNA replication by a fork approaching from the opposite direction. A previous study in *Xenopus* extracts has demonstrated that DHX36, a 3’-5’ helicase, can promote replisome bypass of a leading strand G4 by unwinding DNA downstream (5’) of the G4 (*6*). However, this mechanism requires DHX36 binding to the G4 prior to replisome arrival, which is incompatible with a fork restart function due to the sequestration of G4s inside stalled replisomes. Alternatively, accessory DNA helicases translocating in 5’-3’ direction on the lagging strand template may promote replisome bypass of a G4 analogous to RTEL1-mediated replisome bypass of DNA-protein crosslinks (DPCs) (*58*). Consistent with this possibility, the 5’-3’ DNA helicases Pif1 in budding yeast (*59, 60*) and FANCJ in human cells (*7, 61, 62*) have been demonstrated to promote the replication of G4 DNA. However, whether these helicases can promote the reactivation of stalled forks remains to be determined. Approaches established here will be useful to address this question.

Unexpectedly, our analysis of the CMG stall mechanism has yielded evidence in support of a novel inchworm mechanism for CMG translocation that deviates from rotary hand-over-hand mechanisms suggested by current models (*30, 36*). The translocation can be described as follows (**Figure 7**): Starting from the planar state **(i)**, Mcm6, -2, -5 and -3 coordinate DNA at nucleotide positions 1+2, 3+4, 5+6 and 7+8, respectively, in 3’-5’ orientation. Mcm7 and -4 do not contact the DNA, leaving positions 9+10 and 11+12 unoccupied. Transition into the spiral state **(ii)** involves the disruption of the Mcm2/5 PS1-H2 interface and movement of the Mcm6 and -2 PS1/H2I loops towards the 5’ proximal end of the DNA to engage nucleotides 13+14 and 15+16, respectively, while Mcm5 and -3 remain bound to the DNA on the 3’ proximal end of the spiral. Moreover, the planar-to-spiral rearrangement allows the Mcm7 and -4 PS1/H2I loops to engage base positions 9+10 and 11+12, respectively. Sequential binding of the Mcm7, -4, -6, and -2 PS1/H2I loops to DNA likely determines the step sizes of MCM subunits during the planar-to-spiral transition. Notably, the 12 nt steps for Mcm2 and -6 between states (i) and (ii) translate into an 8 nt 3’-5’ advance by the CMG, which coordinates nucleotide position 8 via Mcm3 in planar state (i) and position 16 via Mcm2 in spiral state (ii) at the 5’ proximal end of the DNA. Next, Mcm5, -3, -7, and -4 sequentially dissociate from nucleotide positions 5-12 and move in 3’ to 5’ direction to reestablish the planar conformation. An intermediate stage of this rearrangement, captured in G4 stall state 2, involves the dissociation of nucleotide positions 5-9 from the Mcm5, -3, and -7 PS1/H2I loops and the approach of the Mcm2 PS1/H2I loops by those of Mcm5 **(iii)**. Completion of the cycle involves the restoration of the Mcm2/5 PS1-H2 interface and engagement of nucleotide positions 17+18 and 19+20 by Mcm5 and -3, respectively, whereas the Mcm7 and -4 PS1/H2I loops remain dissociated from DNA **(iv)**. The Mcm6 and -2 PS1/H2I loops remain bound at positions 13+14 and 15+16, respectively, throughout this transition. Thus, the 12 nt translocation of the Mcm5 and -3 PS1/H2I loops between spiral and planar states corresponds to a 4 nt 3’-5’ advance by the CMG, from nucleotide position 16, coordinated by Mcm2 in spiral state (ii), to nucleotide position 20, coordinated by Mcm3 in the planar state (iv). Collectively, our model thus suggests that CMG traverses 12 nucleotides in one translocation cycle via sequential 8- and 4-nucleotide steps.

**Figure 7:**
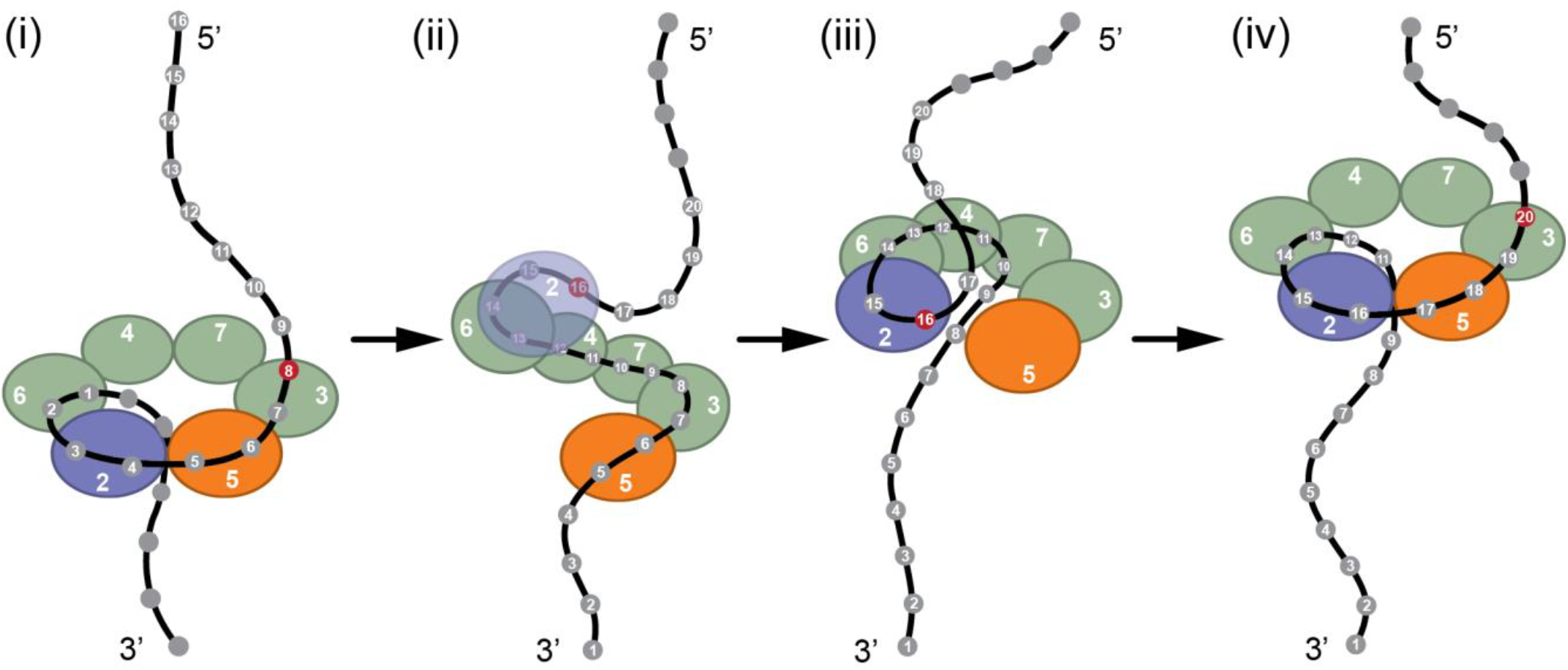
Translocation model. PS1/H2I loops of Mcm2 (blue), Mcm5 (orange), and Mcm6/4/7/3 (green) are depicted as spheres. DNA nucleotide positions are numbered in 3’-5’ orientation with “1” marking the 3’ proximal end of MCM-DNA contacts at the beginning of the translocation cycle. DNA positions highlighted in red mark the 5’ proximal end of MCM-DNA contacts in the respective states.

We suggest that the restructuring of the Mcm2/5 interface is in part mediated by ATP turnover in the Mcm5 ATP-binding site, which is supported with mutational studies demonstrating a critical role for conserved active site residues in Mcm5 for CMG helicase activity (*37*). In our model, ATP-binding to Mcm5 establishes the Mcm2/5 PS1-H2 interface, while ATP-hydrolysis or product release would disengage Mcm2 PS1 from Mcm5 H2. This is consistent with the observation that Mcm5 is occupied by AMP-PNP or ATP in the planar states of CMG bound to normal forks or stalled at a G4, respectively, but nucleotide-free in the spiral state of normal fork conformation 2 (*29*). Mutational analyses demonstrate that ATP turnover by Mcm3 and -7, which follow Mcm5 in the MCM ring, also drive CMG helicase activity, while the remaining ATPase sites are not essential for DNA unwinding (*30, 37*). We hypothesize that ATP turnover at these ATPase sites that are clustered on one side of the Mcm2/5 interface drive the asymmetric structural rearrangements associated with the planar-to-spiral transitions proposed by us.

The CMG inchworm mechanism introduced here is unprecedented among hexameric helicases, which are thought to operate either by rotary or concerted mechanisms (*34, 35*). However, we note that our model bears similarity to the earlier “pump-jack” model (*38*). Moreover, our inchworm model is analogous to the helical inchworm model proposed for the phi29 genome packaging motor, a distantly related homopentameric ATPase (*63, 64*). Based on this analogy, we predict that DNA translocation by CMG occurs in bursts, involving successive 8 and 4 nt steps and likely consuming fewer than 6 ATP molecules per cycle. Careful kinetic analyses will be required to validate these predictions. Furthermore, we expect that approaches established here will facilitate the capture of additional intermediate CMG translocation states in future structural studies.

## Acknowledgments

We thank M. Jason de la Cruz at the MSKCC Richard Rifkind Center for cryo-EM for assistance with data collection and the MSKCC HPC group for assistance with data processing and Lucia Wang for help with artwork.

## Author contributions

S.B. and B.A. performed experiments and analyzed the data. C.K. performed yeast replication experiments. S.D., J.G.B., S.B. and K.M. purified human replication proteins. D.R. and R.H. supervised the project and analyzed the data.

## Supplementary Materials

Materials and Methods

Figures S1-S12

Tables S1-S7

