## Supplemental methods and figures for "Cryo-EM structures of G4-stalled CMG reveal inchworm mechanism of DNA translocation"

**Supplementary Materials**

**Materials and Methods**

***Protein purification***

*Yeast proteins*

Yeast replication proteins were purified as described previously (*5, 44, 65, 66*).

*Human proteins*

hCMG, hPol ε, AND-1, CLASPIN, TIM-TIPIN, hPol δ and hPol α were purified from insect cells using the biGBac baculovirus expression system (*67*). Hi5 cells were infected at a density of $2.5 \times 10^6$ cells per mL and incubated for 72 h (unless indicated otherwise) post infection.

hRFC and CTF18-RFC were purified after overexpression in budding yeast cells. hPCNA and hRPA were purified after overexpression in *E. coli*. Open reading frames (ORFs) were codon-optimized for expression in the respective expression system. For expression in yeast, proteins were expressed from galactose inducible expression vectors stably integrated into the genome. 12 L of cells were grown at 30 °C in YP-GL (YP + 2% glycerol + 2% lactic acid) to a density of $2\text{-}4 \times 10^6$ cells/mL. Galactose was then added to a final concentration of 2% and incubation was continued for an additional 4 hours before harvesting the cells by centrifugation. The cell pellet was washed once with 200 mL of ice cold 1 M sorbitol /25 mM HEPES-KOH pH 7.6 followed by a wash with 50 mL of respective lysis buffer. Cells were then resuspended in half of the volume of lysis buffer and frozen dropwise in liquid nitrogen. Frozen cell popcorn was stored at -80 °C until further processing. Frozen popcorn was crushed in a freezer mill for six 2-min cycles at a rate of 15 impacts per second. Crushed cell lysate was clarified by centrifugation at 42,000 rpm for 45 min (T647.5 rotor). Purified proteins were snap frozen in liquid nitrogen and stored at -80°C.

*hCMG purification*

The cell pellet obtained from 1 L of Hi5 cells was resuspended in 200 mL of hCMG lysis buffer (45 mM HEPES-KOH pH 7.6/0.02% NP40-S/10% glycerol/300 mM KCl/1 mM DTT) supplemented with protease inhibitor cocktail (Thermo Scientific). Once resuspended, cells were lysed via sonication for 2 min (12 cycles of 10 s on, 20 s off) and the lysate was clarified by centrifugation at 42,000 rpm (T647.5 rotor) for 30 min. The soluble phase was incubated with 1 mL of pre-equilibrated M2 agarose anti-FLAG beads at 4°C for 2 h. The beads were washed with 30 column volumes (CV) of hCMG lysis buffer. Bead-bound protein was eluted with 1 CV of hCMG lysis buffer supplemented with 0.5 mg/mL FLAG peptide via gravity flow followed by 2 CV supplemented with 0.25 mg/mL FLAG peptide. FLAG eluates were pooled and incubated with 0.5 mL Strep-Tactin Superflow beads at 4°C for 1 h. The beads were washed with 20 CV of lysis buffer and the bound proteins were eluted with 8 x 1 CV of hCMG lysis buffer supplemented with 30 mM biotin.

Peak fractions containing protein were pooled and diluted with an equal volume of hCMG lysis buffer without KCl prior to fractionation on MiniQ PC 3.2/3 column with a salt gradient of 150 – 1000 mM KCl over 30 CV. Peak fractions from the MonoQ step were pooled and dialyzed against CMG storage buffer (45 mM HEPES-KOH pH 7.6/0.02% NP40-S/10% glycerol/ 200 mM potassium acetate, 1 mM DTT). hCMG yield was around 0.3 mg per L of insect cell culture.

*hPol ε purification*

The cell pellet from 0.5 L of infected Hi5 culture was resuspended in 100 mL of Pol ε lysis buffer (40 mM HEPES-KOH pH 7.6/0.02% NP40-S/10% glycerol/100 mM NaCl/1 mM DTT) supplemented with protease inhibitor cocktail (Thermo Scientific). Cells were lysed via sonication for 2 min (12 cycles of 10 s on, 20 s off) and the lysate was clarified by centrifugation at 42,000 rpm (T647.5 rotor) for 30 min. The soluble extract was incubated with 1 mL of with pre-equilibrated M2 agarose anti-FLAG beads at 4°C for 3 h. The beads were washed with 30 CV of Pol ε lysis buffer. Bead-bound protein was eluted with 1 CV of Pol e lysis buffer supplemented with 0.5 mg/mL FLAG peptide via gravity flow followed by 2 CV supplemented with 0.25 mg/mL FLAG peptide. FLAG eluates were pooled and fractionated on a MonoQ 5/50 column with a salt gradient of 150 – 600 mM NaCl over 20 CV. Peak fractions from the MonoQ step were pooled and spin concentrated (Amicon Ultra 30K centrifugal filters) prior to fractionation on a Superdex 200 10/300 size exclusion column pre-equilibrated in Pol ε storage buffer (40 mM HEPES-KOH pH 7.6/0.02% NP40-S/10% glycerol/200 mM potassium acetate/1 mM DTT). The final yield was around 0.28 mg of protein from 0.5 L of insect cell culture.

*AND-1 purification*

The cell pellet from 0.5 L of infected Hi5 culture was resuspended in 100 mL of AND-1 lysis buffer (25 mM HEPES-KOH pH 7.6/0.02% NP40-S/10% Glycerol/300 mM NaCl/1 mM DTT) supplemented with protease inhibitor cocktail (Thermo Scientific). Cells were lysed by sonication for 2 min (12 cycles of 10 s on, 20 s off) and the lysate was clarified by centrifugation at 42,000 rpm (T647.5 rotor) for 30 min. The soluble extract was incubated with 0.5 mL of pre-equilibrated M2 agarose anti-FLAG beads at 4°C for 2 hours. The beads were washed with 20 CV of AND-1 lysis buffer. Bead-bound protein was eluted with 1 CV of AND-1 lysis buffer supplemented with 0.5 mg/mL FLAG peptide via gravity flow followed by 2 CV supplemented with 0.25 mg/mL FLAG peptide. FLAG eluates were pooled and diluted with an equal volume of AND-1 lysis buffer without NaCl and fractionated on a MonoQ 5/50 column using a salt gradient of 150 – 1000 mM NaCl over 20 CV. Peak fractions from the MonoQ step were pooled and concentrated (Amicon Ultra 30K centrifugal filters) prior to fractionation on a Superdex 200 10/300 column pre-equilibrated with AND-1 storage buffer (25 mM HEPES-KOH pH 7.6/0.02% NP40-S/10% glycerol/200 mM potassium acetate/1 mM DTT). The yield was around 0.15 mg protein from 0.5 L of insect cell culture.

*CLASPIN purification*

0.5 L of Hi5 cells at a density of $2.5 \times 10^6$ cells per mL were infected with CLASPIN baculovirus and the cells were collected 66 h after infection. The cell pellet was resuspended in 100 mL of CLASPIN lysis buffer (25 mM HEPES-KOH pH-7.2/150 mM KCl/5 % glycerol/0.01 % NP40S/1 mM DTT) supplemented with protease inhibitor cocktail (Thermo Scientific). The cells were lysed via sonication for 2 min (12 cycles of 10 s on, 20 s off) and the lysate was clarified by centrifugation at 42,000 rpm (T647.5 rotor) for 30 min. The soluble extract was incubated with 0.5 mL of pre-equilibrated Strep-Tactin Superflow beads at 4°C for 3 hours. The beads were washed with 40 CV of CLASPIN lysis buffer. Bead-bound protein was eluted with 8 CV of CLASPIN lysis buffer supplemented with 30 mM biotin via gravity flow. Eluates were pooled and concentrated (Amicon Ultra 30K centrifugal filters) prior to fractionation on a Superose 6 10/300 gel filtration column pre-equilibrated in CLASPIN storage buffer (25 mM HEPES-KOH pH 7.6/0.02% NP40-S/10% glycerol/200 mM potassium acetate/1 mM DTT). The protein yield was around 0.1 mg per 0.5 L of insect cell culture.

*TIM-TIPIN purification*

50 mL of Hi5 cells were infected at a density of $2.5 \times 10^6$ per mL and collected 72 hours post-infection and pelleted by centrifugation at 3000 rpm in a swinging bucket rotor at 4°C. The cell pellet was resuspended in 20 mL of TIM-TIPIN lysis buffer (25 mM HEPES-KOH pH 7.2/0.01% NP40-S/5% Glycerol/150 mM KCl/1 mM DTT) supplemented with protease inhibitor cocktail (Thermo Scientific). Cells were lysed via ultra-sonication pulses of 10 s on and 30 s off for 12 cycles and the lysate was clarified by centrifugation at 42,000 rpm (T647.5 rotor) for 30 min. The soluble extract was passed over 400 μL of pre-equilibrated M2 agarose anti-FLAG bead at 4°C. The beads were washed with 40 CV of TIM-TIPIN lysis buffer. Bead-bound protein was eluted with 2 CV of TIM-TIPIN lysis buffer supplemented with 0.2 mg/mL FLAG peptide and mixing on hot-dog roller for 15 min at 4°C, followed by 2 CV supplemented with 0.1 mg/mL FLAG peptide. FLAG eluates were pooled and applied onto a 1 mL MonoQ column equilibrated in TIM-TIPIN lysis buffer. Bound proteins were eluted with a salt gradient of 150 – 1000 mM KCl applied over 20 CV. Peak fractions from the MonoQ column were pooled and concentrated using Amicon Ultra-0.5 (30 kDa cutoff) filter (Millipore-Merck) prior to fractionating on a 24 mL Superdex 200 size exclusion column pre-equilibrated with 25 mM Tris pH-7.2/150 mM NaCl/5 % glycerol/0.01 % NP40S/1 mM DTT. The peak fractions were concentrated on an Amicon Ultra-0.5 (30 kDa cutoff) filter (Millipore-Merck) before storage at -80 °C. The yield of purified TIM-TIPIN complex was around 0.31 mg per 50 mL of insect cell culture.

*hPol $\delta$ purification*

The cell pellet from 1 L of infected Hi5 cells was resuspended in 200 mL of Pol $\delta$ lysis buffer (30 mM Tris HCl pH 7.5/300 mM NaCl/0.03% NP-40S/6% glycerol/1 mM DTT) supplemented with protease inhibitor cocktail (Thermo Scientific). Cells were lysed via sonication for 2 min (12 cycles of 10 s on, 20 s off) and the lysate was clarified by centrifugation at 42,000 rpm (T647.5 rotor) for 30 min. The soluble extract was incubated with 1 mL of pre-equilibrated Strep-Tactin Superflow beads at 4°C for 3 hours. The beads were washed with 40 CV of Pol $\delta$ lysis buffer. Bead-bound protein was eluted with 8 CV of Pol $\delta$ lysis buffer supplemented with 30 mM biotin via gravity flow. The eluates were pooled and diluted to bring the NaCl concentration to 100 mM and supplemented with 0.5 mM ATP and 5 mM magnesium chloride, and incubated for 30 minutes at 4°C. The eluates were pooled and diluted to bring the NaCl concentration to 100 mM, supplemented with 0.5 mM ATP and 5 mM magnesium chloride, and incubated for 30 min at 4°C. Subsequently, the eluate was applied onto a 1 mL Capto HiResQ column equilibrated in 50 mM Tris pH-8.0/10 % glycerol/0.01 % NP40-S/100 mM NaCl/1 mM DTT. Bound protein was eluted by applying a gradient of 100 – 1000 mM

NaCl over 30 CV. Peak fractions were pooled, supplemented with 0.5 mM ATP and 5 mM magnesium chloride, incubated for 30 minutes at 4°C, concentrated in a spin concentrator, and fractionated on a 24 mL Superdex 200 size exclusion column pre-equilibrated with 25 mM Tris pH-7.2/150 mM NaCl/10 % glycerol/0.005 % Tween 20/1 mM DTT. The peak fractions were concentrated on a spin concentrator before storage at -80 °C. The final yield for hPol $\delta$ was around 0.15 mg of protein per 1 L of insect cell culture.

*hPol $\alpha$ purification*

The cell pellet from 1 L of infected Hi5 cells was resuspended in 200 mL of Pol $\alpha$ lysis buffer (25 mM HEPES-NaOH pH 7.5/10% Glycerol/0.02% NP40-S/300 mM NaCl/1 mM DTT) supplemented with protease inhibitor cocktail (Thermo Scientific). Cells were lysed via sonication for 2 min (12 cycles of 10 s on, 20 s off) and the lysate was clarified by centrifugation at 42,000 rpm (T647.5 rotor) for 30 min. The soluble extract was incubated with 1 mL of pre-equilibrated Strep-Tactin Superflow beads for 4 h at 4°C. The beads were washed extensively with 30 CV of Pol $\alpha$ lysis buffer. Bead-bound protein was eluted with 8 CV of Pol $\alpha$ lysis buffer supplemented with 30 mM biotin via gravity flow. Eluates were pooled and diluted gradually with two volumes of Pol $\alpha$ lysis buffer without NaCl and fractionated on a Capto HiResQ 5/50 column using a salt gradient of 100 – 1000 mM NaCl over 20 CV. Peak fractions from the Capto HiResQ 5/50 step were pooled and concentrated (Amicon Ultra 30K centrifugal filters) prior to fractionation on a Superdex 200 10/300 size exclusion column pre-equilibrated in Pol a storage buffer (25 mM HEPES-NaOH pH 7.5/10% glycerol/0.02% NP40-S/200 mM potassium acetate/1 mM DTT). The final yield was around 0.15 mg of purified hPol $\alpha$ per 1 L of insect cell culture.

*hRFC purification*

Crushed cell powder was thawed and an equal volume of RFC lysis buffer (25 mM HEPES-KOH pH7.6/20% sucrose/5 mM magnesium acetate/0.02% NP40-S/0.02% Triton X-100/0.5% inositol/1 mM ATP/300 mM NaCl/1 mM DTT), supplemented with protease inhibitor cocktail (Thermo Scientific), was added to resuspend the lysate. The cell lysate was clarified by centrifugation at 40,000 rpm (T647.5 rotor) for 45 min and the soluble extract was incubated with 1 mL of pre-equilibrated M2 agarose anti-FLAG beads for 2 hours at 4°C for 2 hours. The beads were washed extensively with 15 CV of RFC lysis buffer. Bead-bound protein was eluted with 1 CV of RFC lysis buffer supplemented with 0.5 mg/mL FLAG peptide via rotation on hot dog roller for 10 min) followed by 2 CV supplemented with 0.25 mg/mL FLAG peptide. Eluates were pooled and the FLAG affinity tag was cleaved by incubation with 10-fold molar excess of TEV protease overnight at 4°C. The salt concentration of the protein sample was lowered to 200 mM NaCl by gradually adding RFC lysis buffer without NaCl prior to fractionation on a MonoS 5/50 column using a salt gradient of 200 – 1000 mM NaCl over 20 CV. Peak fractions were pooled and dialyzed against RFC lysis buffer containing 200 mM NaCl and fractionated on a HiTrap Heparin HP(1 mL) column using a gradient of 200- 1000 mM NaCl over 20 CV. Peak fractions were pooled and dialyzed against RFC storage buffer (25 mM HEPES-KOH pH7.6/20% sucrose/5 mM magnesium acetate/0.02% NP40-S/0.02% Triton X-100/0.5% inositol/1 mM ATP/300 mM potassium acetate/1 mM DTT) prior to storage. The protein yield was around 1 mg of purified hRFC per 12L of yeast cells.

*CTF18-RFC purification*

Crushed cell powder was thawed and an equal volume of CTF18 lysis buffer (50 mM HEPES-KOH pH7.6/10% glycerol/0.02% NP40-S/1 mM EDTA/1 mM EGTA/150 mM KCl/1 mM DTT) , supplemented with protease inhibitor cocktail (Thermo Scientific), was added to resuspend the lysate to homogeneity. The cell lysate was clarified by centrifugation at 40,000 rpm (T647.5 rotor) for 30 min and the soluble extract was passed through 1 mL of pre-equilibrated M2 agarose anti-FLAG beads at 4°C using an Econo Pump (Bio-Rad). The beads were washed extensively with 15 CV of RFC lysis buffer. Bead-bound protein was eluted with 3 CV of CTF18 lysis buffer supplemented with 0.5 mg/mL FLAG peptide (via rotation on hot dog roller for 10 min) followed by 2 CV of lysis buffer. Eluates were pooled and fractionated on a MonoQ 5/50 column using a salt gradient of 150 – 1000 mM KCl over 20 CV. Peak fraction from the MonoQ step was fractionated on a Superdex 200 10/300 size exclusion column pre-equilibrated with CTF18 storage buffer (25 mM HEPES-KOH pH 7.6/0.02% NP40-S/10% glycerol/1 mM EDTA/1 mM EGTA/300 mM potassium acetate/1 mM DTT). The final yield for CTF18-RFC was around 2.5 mg of protein per 12 L of yeast cells.

*hPCNA purification*

BL21-Codonplus DE3 (RIL) cells (Agilent Technologies) were freshly transformed with the PCNA expression construct. 1 L of cells were grown at 37°C to $OD_{600}$ = 0.6, cooled on ice for 20 min and ethanol was added to a final concentration of 2%. Expression was induced by addition of 1 mM IPTG and incubation continued at 37°C for 3 h. Cells were harvested by centrifugation at 4000 rpm for 15 min and the cell pellet was stored at -80°C until further processing. The cell pellet was thawed and resuspended in 50 mL of PCNA lysis buffer (50 mM sodium phosphate pH 7.6/400 nM NaCl/10 mM imidazole/1 mM DTT) supplemented with protease inhibitor cocktail (Thermo Scientific) and 0.1 mg/mL of lysozyme. Cells were lysed via sonication 1.5 min (3 cycles of 30 s on, 90 s off) and the lysate was clarified by centrifugation for 30 min at 12,000 rpm (SS34 rotor). The soluble phase was passed over 2 mL of equilibrated $Ni^{2+}$-NTA agarose resin via gravity flow in a disposable column. Beads were washed with 15 CV of PCNA wash buffer (25 mM Tris-HCl pH 7.5/1 mM EDTA/150 mM NaCl/0.02% NP40-S/10% glycerol/10 mM imidazole/1 mM DTT). Bead-bound protein was eluted with 6 CV of PCNA elution buffer (25 mM Tris-HCl pH 7.5/1 mM EDTA/150 mM NaCl/0.02% NP40-S/10% glycerol/200 mM imidazole/1 mM DTT). Eluates were pooled and fractionated on a MonoQ 5/50 column using a salt gradient of 150 – 1000 mM NaCl over 20 CV. Peak fraction from the MonoQ step was fractionated on a Superdex 200 10/300 size exclusion column pre-equilibrated in PCNA storage buffer (25 mM Tris-HCl pH 7.5/1 mM EDTA/300 mM potassium acetate/0.02% NP40-S/10% glycerol/10 mM imidazole/1 mM DTT). The protein yield was around 8 mg of purified hPCNA per 1 L of cells.

*hRPA purification*

The RPA expression plasmid, p11d-tRPA, was a gift from Marc Wold (University of Iowa). RPA was purified as reported previously (*68*). 8 L of BL21-Codonplus DE3 (RIL) transformed with RPA expression plasmid were grown at 37 °C to $OD_{600}$ = 0.8. RPA expression was induced by the addition of 1 mM IPTG and continued incubation for 3 h. Cells were harvested and resuspended in 130 mL of 30 mM HEPES-KOH pH 7.5/5 mM DTT/1 mM EDTA/0.5% inositol/1 mM

PMSF/0.02% Tween 20/50 mM KCl and protease inhibitor cocktail. Cells were lysed by three passages through a French Press at 20,000 psi and the lysate was cleared by centrifugation in a T647.5 rotor at 30,000 rpm for 40 min. One-third of the supernatant (fraction 1) was diluted with 2.5 volumes of Buffer J (30 mM HEPES-KOH pH 7.5/5 mM DTT/1 mM EDTA/0.5% inositol/0.02% Tween 20/10% glycerol) + 50 mM KCl and loaded onto a 100 mL Affi-Gel Blue column equilibrated in the same buffer. The column was washed with 3 CV equilibration buffer, 3.5 CV Buffer J + 900 mM KCl, and 3 CV Buffer J + 50 mM KCl and 0.5 M NaSCN. Protein (fraction 2) was eluted with Buffer J + 1.5 M NaSCN. Fraction 2 was loaded onto an 11 mL column of hydroxyapatite (BioRad MacroPrep) equilibrated with Buffer J + 50 mM KCl and the column was washed with 2 CV of equilibration buffer, 2 CV of Buffer J + 800 mM KCl, and then 2 CV of equilibration buffer. Protein (fraction 3) was eluted with Buffer J + 50 mM KCl and 100 mM potassium phosphate (pH 7.5). Fraction 3 was diluted with Buffer J + 30 mM KCl to a conductivity equivalent to Buffer J + 100 mM KCl and loaded onto a 1 mL Mono-Q column equilibrated in Buffer J + 50 mM KCl. The column was washed with 4 CV of equilibration buffer and 4 CV Buffer J + 100 mM KCl. Protein was eluted with a 20 CV gradient of 200 to 400 mM KCl in Buffer J. hRPA (fraction 4) was pooled based on SDS-PAGE. Fraction 4 (3.3 mg/ml, 2.8 mL) was aliquoted, frozen in liquid $N_2$, and stored at -80 °C.

***DNA templates***

*DNA templates for use in the reconstituted yeast DNA replication system*

DNA templates were generated by ligating small oligonucleotide inserts containing a pre-formed G4 structure into the ARS305-containing plasmid p1400. p1400 was linearized by digestion with BbsI and XcmI to generate non-complementary overhangs. The linearized plasmid DNA was isolated by fractionation on a 10-40 % sucrose gradient (36 mL volume) prepared in 20 mM Tris pH-7.6/1 M NaCl/1 mM EDTA. Gradients were centrifuged for 20 h at 27,000 rpm and 20 ºC in an AH-629 swinging bucket rotor (Thermo Scientific). Peak fractions corresponding to the linearized form of p1400 were concentrated and buffer-exchanged to 10 mM Tris pH-8.0/1 mM EDTA (1xTE) using Amicon Ultra-15 centrifugal filters (10 kDa cutoff; Millipore-Merck).

Inserts harboring a pre-formed G4 were generated by annealing PAGE purified, 5'-phosphorylated oligonucleotides (**Table S1**). Oligonucleotides were electrophoresed on 10 % urea polyacraylamide gels and extracted using the crush-and-soak method (*46*). Complementary oligonucleotides (10 µM each) were mixed in 500 µL of 20 mM Tris pH-8.0/ 25 mM potassium chloride/10 mM magnesium chloride and annealed by heating to 95 ºC for 5 min, followed by cooling to 10 ºC at a rate of 1 ºC/minute in a thermocycler. Annealed products were electrophoresed on 8 % (39:1 acrylamide:bis-acrylamide) native polyacrylamide gels, excised from the gel and eluted from the gel slices by soaking overnight at 4 ºC in 10 mM Tris pH-8.0/50 mM potassium acetate/1 mM EDTA. Ligations were performed similarly as described previously (*69*). Each ligation reaction was carried out in a volume of 80 mL containing 0.5 nM linearized p1400, 1.5 nM purified insert and 50 mM Tris pH-7.6/10 mM magnesium chloride/2.5 mg/mL BSA/0.1 mM ATP/1 mM DTT and 8000 units of T4 DNA ligase. Reactions were incubated overnight at 16 ºC. Subsequently, ligation reactions were concentrated using Amicon Ultra-15 filter (Millipore-Merck). Re-circularized supercoiled plasmid from each ligation mixture was separated using cesium chloride ultracentrifugation. For this, concentrated ligation reactions were reconstituted into 0.8 g/mL cesium chloride using a cesium chloride solution in 1xTE and ultracentrifuged in a TV-1665

(Thermo Scientific) vertical rotor at 60,000 rpm for 16 h at 4 ºC. The supercoiled plasmid DNA band was recovered by aspiration, extracted 3 times with butanol, concentrated and buffer-exchanged into 1xTE using Amicon Ultra-0.5 (3 kDa cutoff) filters (Millipore-Merck).

*DNA templates for use in reconstituted human DNA replication system*

Forked DNA templates for human replication assays were designed similarly as described previously (*70*). The templates were prepared in two steps: 1) Preparation and isolation of circularized plasmid with G4 inserts similar to the yeast replication templates. 2) Linearization of the circular plasmid to asymmetrically position the G4 structures between the termini, followed by ligation of forked DNA to one end. BstXI and AfeI sites were introduced into p1400 to serve as sites of linearization and fork ligation. This modified vector, p1432, was first linearized with BbsI and XcmI, followed by ligation of a G4 insert (**Table S1**) as described above for the yeast replication templates. Purified supercoiled plasmids with pre-formed G4 structures were subsequently linearized with BstXI and AfeI. These linearized G4 fragments with one cohesive end were purified by 10-40 % sucrose gradient centrifugation (2.2 mL tubes) in 20 mM Tris pH-7.6/1 M NaCl/1 mM EDTA. Centrifugation was carried out at 53,000 rpm for 4 hours at 20 ºC in a TLS55 swinging bucket rotor (Beckman Coulter). Peak fractions corresponding to the linearized G4-containing vector DNA were concentrated and buffer-exchanged to 10 mM Tris pH-8.0/1 mM EDTA (1xTE) using Amicon Ultra-0.5 (3 kDa cutoff) centrifugal filters (Millipore-Merck). The isolated linear vector DNA was ligated to a forked DNA molecule with complementary overhangs to BstXI site, prepared by annealing PAGE-purified oligonucleotides (**Table S2**). 100-150 mL ligation reactions were performed overnight with 2,000 units of T4 DNA ligase (NEB) at 16 ºC in NEBuffer r2.1 (NEB) supplemented with 1 mM ATP. Reactions included 5-10 pmol of linearized G4-containing vector DNA and a 10-fold molar excess of fork DNA. Subsequently, T4 DNA ligase was heat-inactivated at 65 ºC for 10 min and ligation reactions were treated with 30 units of AfeI to monomerize fork dimers blunt-ligated at their distal ends. Monomerized fork templates were purified again on 10-40 % sucrose gradients (2.2 mL tubes), centrifuged for 4 hours at 53,000 rpm and 20 ºC in a TLS55 swinging bucket rotor (Beckman Coulter). Fractions corresponding to monomeric forked DNA templates were pooled, concentrated and buffer exchanged to 1xTE using Amicon Ultra-0.5 (3 kDa cutoff) filter (Millipore-Merck).

*DNA substrates for helicase and 3' exonuclease protection assays*

Templates were prepared by annealing PAGE purified oligonucleotides (TABLE 2) with one of the complementary oligonucleotides being 5' radiolabeled. The G4-containing tracking strand was radiolabeled for the 3' exonuclease protection assay, whereas the non-G4 strand was radiolabeled for the helicase assays. Oligonucleotide (0.25 μM) was radiolabeled in a 15 μL reaction containing 0.5 μM [$\gamma$-$^{32}$P]ATP, 10 units of T4 PNK (NEB) for 1 h at 37 ºC, followed by heat inactivation at 80 ºC for 20 min. Reactions were supplemented with 50 mM potassium acetate and 1.5 times molar excess of complementary oligonucleotide (**Table S3**). Annealing was carried by heating the mixture to 95 ºC for 5 min, followed by cooling to 10 ºC at a rate of 1 ºC/min using a thermocycler. Annealed products were separated on 8 % (39:1 acrylamide:bisacrylamide) native polyacrylamide gels, excised from the gel and eluted from the gel slices by soaking overnight in 10 mM Tris pH-8.0/50 mM potassium acetate/1 mM EDTA.

*DNA template for 5' exonuclease protection assay*

The tracking strand was prepared by RNA splint-ligation of two oligonucleotides and subsequently annealed to the complementary non-tracking strand. All oligonucleotides, except the RNA oligonucleotide, were PAGE purified as described above. A 43 nt long oligonucleotide (2970) that corresponds to the 3' portion of the tracking strand is first radiolabeled at its 5' end. For this, oligonucleotide (1 μM) is radiolabeled in a 30 μL reaction containing 0.5 μM γ-[$^{32}$P]-ATP and 10 units of T4 PNK (NEB) for 1 hour at 37 ºC. The reaction is subsequently supplemented with 25 μM ATP and incubated for another 30 minutes to phosphorylate the remaining 5' ends, followed by heat inactivation of T4 PNK. The reaction mix is then supplemented with 10 mM potassium chloride, 1.5 times molar excess of a second oligonucleotide (2990) that corresponds to the 5' part of the tracking strand and 2.5 times molar excess of a RNA oligonucleotide (2973). The three oligonucleotides were annealed, resulting in an RNA-DNA hybrid juxtaposing the 5' labeled phosphate of 2970 to the 3' end of 2290. This juxtaposition of 2990 and 2970 is equivalent to full-length tracking strand with a nick opposite the complementary RNA oligonucleotide that was sealed by addition of 50 units of SplintR ligase (NEB) and incubation for 4 h at 25 ºC. The RNA oligonucleotide was subsequently degraded by addition of 50 pmoles of yeast RNase H1 and 20 μg RNase A (Themo Scientific) at 30 ºC for 1 hour. The final reaction volume was 40 μL. The complementary non-tracking strand (2884) was added to the reaction mix containing and annealed by heating the reaction to 95 ºC and slow cooling to 10 ºC at a rate of 1 ºC/min in a thermocycler. The reaction mixture was supplemented with 0.5 % SDS and treated with 0.8 units of Proteinase K (NEB) for 1 hour at 37 ºC. Templates were separated on 8 % (39:1 acrylamide:bisacrylamide) native polyacrylamide gels, excised from the gel and eluted from the gel slices by soaking the gel slices overnight in 10 mM Tris pH-8.0/50 mM potassium acetate/1 mM EDTA.

*DNA template for cryo-EM analysis*

Templates were prepared by annealing PAGE purified oligonucleotides and extracting the annealed products from native polyacrylamide gels. Complementary oligonucleotides (2883 and 2884, 20 mM each) were mixed in 500 μL of 20 mM Tris pH-8.0/25 mM potassium chloride/10 mM magnesium chloride and annealed by heating to 95 ºC for 5 min, followed by cooling to 10 ºC at a rate of 1 ºC/min in a thermocycler. Annealed products were electrophoresed on 8 % (39:1 acrylamide:bisacrylamide) native polyacrylamide gels, excised from the gel and eluted from the gel slices by soaking the gel slices overnight in 10 mM Tris pH-8.0/50 mM potassium acetate/1 mM EDTA. The DNA was concentrated to a final concentration of 50 - 100 μM using an Amicon Ultra-0.5 (3 kDa cutoff) filter (Millipore-Merck).

***3' exonuclease protection assay***

Fifty microliter reactions containing 4 nM template DNA, 20 nM CMG, 20 nM Csm3-Tof1 and 20 nM Mrc1 were assembled in helicase buffer (20 mM HEPES-KOH pH 7.6/100 mM potassium acetate/10 mM magnesium acetate/0.1 mg/mL BSA/2.5 mM DTT) and were incubated with 0.1 mM AMP-PNP for 30 min at 30 ºC. 25 units of Exonuclease T (NEB) were added to the reaction and incubation continued for another 5 min. A 10 μL aliquot that was withdrawn and added to 90 μL of stop buffer (0.5 % SDS/20 mM EDTA/100 mM sodium chloride/125 μg/mL tRNA) served as the 0-minute time point. The remaining reaction was mixed with an equal volume of 10 mM ATP in helicase buffer and 20 μL aliquots were withdrawn at 2, 10, 30 and 50 minute incubation times and reactions stopped by addition of 80 μL of stop buffer followed by extraction with Tris-saturated phenol/chloroform solution (Acros). Aqueous phase contents were ethanol precipitated, rinsed with 70 % ethanol and dried in a rotary dry-vac. The pellets were resuspended in 12 mL of 33 mM NaOH/50 % (v/v) formamide, heated to 95 ºC for 5 min and snap-chilled on ice. Reaction products were electrophoresed on 6 % or 8 % urea polyacrylamide sequencing gels in 1xTBE. Gels were dried on a Whatman sheet, exposed to phosphor-imager and scanned on Typhoon IP scanner (Cytiva).

***5' exonuclease protection assay***

Reactions were carried out in 30 µL of helicase buffer containing 4 nM template DNA and either 20 nM CMG (yeast or human) or a mixture of CMG, Csm3-Tof1/TIM-TIPIN and Mrc1/CLASPIN. The protein/DNA mixtures were incubated for 30 min, either at 30 ºC in the case of the yeast proteins or at 37 ºC in the case of the human proteins. Prior to the addition of ATP, 10 units of T5 exonuclease (NEB) or 22 units of a 5' exonuclease concoction [4 units T7 exonuclease (NEB), 12 units $RecJ_f$ (NEB), 4 units Exonuclease VIII, truncated (NEB), 2 units T5 exonuclease (NEB)] were added to a 20 µL reaction aliquot and incubation continued for 5 min at 30 ºC or 37 ºC, respectively. The exonuclease reaction was stopped by addition of 80 µL of stop buffer. The remaining reaction was mixed with an equal volume of 10 mM ATP in helicase buffer and 20 µL aliquots were withdrawn at designated times and treated with either T5 exonuclease or the 5' exonuclease concoction as above. Reactions were stopped by addition of 80 µL of stop buffer and phenol/chloroform-extracted. Aqueous phase contents were ethanol-precipitated, rinsed with 70 % ethanol and dried in a rotary dry-vac. The pellets were resuspended in 12 µL of 33 mM NaOH/ 50 % (v/v) formamide, heated to 95 ºC for 5 min and snap-chilled on ice. Reaction products were electrophoresed on a 6 % urea polyacrylamide sequencing gel in 1xTBE. Gels were dried onto Whatman paper, exposed to phosphor-imager and scanned on Typhoon IP scanner (Cytiva).

***Helicase assay***

4 nM subtrates were incubated for 30 minutes at 30 ºC with 20 nM of each CMG, Csm3-Tof1 and Mrc1 in a 20 µL reaction volume prepared in helicase buffer. An equal volume of 10 mM ATP in helicase buffer was added to initiate DNA unwinding. At the indicated times, 4 µL aliquots were withdrawn and stopped by mixing with 4 µL of stop buffer containing 0.1 % SDS/40 mM EDTA. Except for the experiments in Supplementary Figure 2, reactions were supplemented with 20 nM un-labeled oligonucleotide corresponding to the labeled strand in template to prevent the re-annealing of the product strands and 40 nM $(dT)_{40}$ (2602) to sequester free CMG, one minute after the initiation of unwinding. Reaction products were electrophoresed on 8 % (39:1 acrylamide:bisacrylamide) native polyacrylamide gels in 1xTAE for 2 hours at 75 V. Gels were dried on a Whatman sheet, exposed to phosphor-imager and scanned on Typhoon IP scanner (Cytiva). The bands were quantified using ImageJ and plotted using GraphPad Prism software.

***Sample preparation for cryo-EM analysis***

0.25 µM yeast or human CMG, Csm3-Tof1/TIM-TIPIN and Mrc1/ CLASPIN were incubated with 5 µM template DNA and 100 µM AMP-PNP for 30 min at 30 ºC (yeast proteins) or 37 ºC (human proteins) in a 20 µL reaction containing 20 mM HEPES-KOH pH 7.6/100 mM potassium acetate/10 mM magnesium acetate/2.5 mM DTT/0.02 % NP40-S. DNA unwinding was initiated by adding 5 mM ATP to the reaction and incubating the reaction for another 30 minutes. 4 µL of a reaction were spotted on graphene oxide layered Quantifoil grids (EM Sciences), incubated for 30 seconds and blotted for 15 or 30 seconds. Grids were plunge-frozen using a Vitrobot II (Thermo Scientific).

**Cryo-EM data collection**

Cryo-EM samples were imaged using a 300keV FEI Titan Krios microscope equipped with a K3 summit direct electron detector (Gatan). Images were recorded with SerialEM (*71*) in super-resolution mode at 29,000x, corresponding to a pixel size of 0.413 Å/px. Movies were recorded over 3 sec at a dose rate of 15 $e^-$/pix/sec (0.05 sec/frame), yielding a total dose of 66 $e^-$/Å$^2$.

**Cryo-EM data processing**

All movies were gain-corrected, 2x Fourier-cropped to 0.826Å/px, CTF estimated, and aligned using whole-frame and local motion correction in CryoSPARC v.4.0.0 (*72*). Blob-based autopicking was used with to select initial particle images within cryoSPARC Live, after which iterative rounds of 2D classification were performed and best 2D class averages were used to generate an *ab initio* 3D model. The initial model was used alongside six false-positive "decoy" 3D noise classes in iterative rounds of heterogenous refinement to select for true positive particles from among the full stack of blob-picked particles. After ten rounds of heterogenous refinement, a further round of 2D classification was used to manually curate a selection of particles for use in training a convolutional neural network particle picker (*73*). All Topaz-picked particles (~1.9M) were extracted at 0.826 Å/px, binned 2x to 1.652 Å/px, and heterogeneously refined as previously described. The final particle stack (294,032) was then re-extracted at 0.826 Å/px prior to Bayesian Polishing in RELION-4 (*74*). Polished particles were imported to CryoSPARC and refined using non-uniform refinement with local CTF estimation and higher-order aberration correction on particles separated by optical group. These particles were 3D classified to distinguish conformational heterogeneity among the MCM C-tier and G-quadruplex, in addition to a population of particles which lacked Tof1-Csm3. These particles were again refined using non-uniform refinement followed by local refinements of each the N-tier, C-tier, Cdc45-GINS, and Tof1-Csm3 with or without signal subtraction of the rest of the complex. Importantly, masks around each domain were 40 Å soft-padded to reduce boundary artifacts, and slightly overlapped with other masks to aid in composite map generation. Individual local refinements were density-modified in Phenix (*75*) and joined into a composite map using Phenix's combine-focus-maps tool.

**Model building and refinement**

Composite maps were used for *de novo* model building using the automated graph neural network ModelAngelo (*76*). The models were manually rebuilt to fit the EM density in COOT (*77*) and geometrically improved using ISOLDE (*78*). Iterative real-space refinement in Phenix and manual correction in COOT was used to refine the model (*79*). All structural figures were generated in PyMOL (Schrodinger, LLC. 2010. The PyMOL Molecular Graphics System, Version 2.5.4) and ChimeraX (*80*).

***DNA replication assays with purified yeast proteins***

Reactions were performed as described (*5*).

***DNA replication assays with purified human proteins***

Reactions were performed at 37 ºC in 25 mM HEPES-KOH pH-7.6/125 mM potassium glutamate/10 mM magnesium acetate/0.1 mg/mL BSA/1 mM DTT. The concentrations of other components in a reaction were as follows: 1 nM DNA template, 0.1 nM hPol δ, 10 nM hPol α, 120 nM hRPA, 20 nM hCMG, 20 nM TIM-TIPIN, 20 nM CLASPIN, 20 nM AND1, 20 nM hPCNA, 20 nM hRFC, 20 nM CTF18-RFC, 20 nM hPol ε, 50 µM AMP-PNP, 40 µM dNTPs, 200 µM G/C/UTP, 5 mM ATP and 33 nM (1 µCi) [$\alpha^{32}$P]dATP. Reactions were performed by pre-incubating hCMG, TIM-TIPIN, CLASPIN and AND1 for 10 min with DNA template in the presence of 0.1 mM AMPPNP. Reactions were then diluted two-fold by mixing an equal volume of reaction mix containing the remaining nucleotide and protein factors. Aliquots were drawn at the indicated times and stopped by either adding 50 mM EDTA for denaturing gel analysis or heat inactivation at 80 ºC for native gel analysis. For denaturing gel analysis, DNA was isolated from the reactions using SpeedBead magnetic carboxylate (Cytiva). For this, 10 µL of stopped reaction mix was added to 18 µL of SpeedBead mix (2 % SpeedBeads/10 mM Tris pH-8.0/2.5 M sodium chloride/20 % PEG 8000/1 mM EDTA). The beads were separated on a magnetic rack and rinsed three times with 85 % (v/v) ethanol. DNA was eluted with 20 µL of 100 mM NaOH and electrophoresed on a 0.8 % agarose gel for 3 hours at 40 V in 30 mM NaOH/2 mM EDTA. Gels were neutralized with 5 % (w/v) TCA, dried onto Whatman paper and exposed to phosphor imager.

For native gel analysis, reactions were heat-inactivated for 20 minutes at 80 ºC and treated with 2.5 units of Quick CIP (NEB) for 30 min to remove phosphate radiolabel from unincorporated nucleotides. The reactions were subsequently supplemented with 0.5 % SDS, 10 mM EDTA and 0.8 units of Proteinase K (NEB) and incubation continued for 30 min. The products electrophoresed for 200 min at 55 V on 0.8 % agarose gels in 1xTAE. Gels were dried on Whatman paper and exposed to phosphor imager.

**Supplementary Figure Legends**

**Figure S1: DNA unwinding by yeast CMG-FPC is inhibited by a G4 specifically in the leading strand template. (A)** Representative CMG-FPC helicase assays. **(B)** Quantitative comparison of CMG and CMG-FPC helicase activities on control and G4-containing DNA templates. Lines mark average of three replicate experiments.

**Figure S2: The block to CMG-FPC progression is independent of G4 topology. (A)** Representative CMG-FPC helicase assays on templates harboring different G4s. Underlined nucleotides indicate G tracts forming G quartets in indicated G4 structures. Red nucleotides indicate G4 loop sequences. **(B)** Quantitation of helicase assays on templates harboring different G4s. Lines mark average of three replicate experiments.

**Figure S3: 3' exonuclease protection assay. (A)** Exo T products on single-stranded vs forked DNA template. Products were separated on sequencing gel and analyzed by phosphor imaging. Schematics on the left indicate the structure of the single-stranded and forked templates. **(B)** CMG-FPC titration experiment. CMG-FPC was assembled onto forked template for 30 minutes in the presence of AMP-PNP, followed by 2 minutes of Exo T digestion. **(C)** Time course analysis of CMG-FPC assembly on forked DNA template. CMG-FPC was assembled onto forked template for indicated time periods, followed by 2 minutes of Exo T digestion. **(D)** Time course analysis of Exo T digestion reaction. CMG-FPC was assembled onto forked DNA template for 30 minutes in the presence of AMP-PNP, followed by the addition of Exo T for indicated time periods.

**Figure S4: The relative stall position of CMG-FPC is independent of the distance of the G4 from the original fork junction. (A)** 3' exonuclease protection assay on DNA template harboring a prototypical G4 in the leading strand template 34 bp downstream of the original fork junction. **(B)** 3' exonuclease protection assay on DNA templates harboring a CEB25 G4 in the leading strand template 34 bp downstream of the original fork junction.

**Figure S5: EM data processing workflow.**

**Figure S6:** Superposition of PDB 2M4P (blue) onto G4 DNA state 1 (grey and red), aligned on the G4.

**Figure S7: Schematic of protein-DNA contacts in G4 stall states 1 and 2.** Black squares: DNA base. Solid black line: DNA backbone ribose. Pink circle: DNA backbone phosphate. Black rounded rectangle: ATP. Grey rounded rectangle: ADP. Circled residues: H2I. Residues in rectangles: PS1. Residues in rhomboid: H2. Dashed black line: Protein-DNA contact.

**Figure S8: (A)** Superposition of DNA in state 1 (green) and state 2 (blue), aligned on the lagging strand template. **(B-D)** Structure of the fork in state 1 (B), state 2 (C) and normal fork conformation 1 (PDB 6SKL; D). Residues that coordinate the DNA and the leading strand base of the last base pair at the fork junction are shown as sticks. Mcm2 has been removed for clarity.

**Figure S9: PS1/H2I loop conformations. (A)** Conformation of the PS1/H2I loop in human CMG-FPC structures. PDB 7PFO (left) corresponds to yeast normal fork conformation 1 and PDB 6XTX (right) corresponds to yeast normal fork conformation 2. Mcm2 is shown in blue, Mcm6, -4, -3, and -7 are shown in green and Mcm5 is shown in orange. **(B)** Intersubunit PS1-H2 interactions, 6SKO **(C)** Close-up of intersubunit PS1-H2 interactions, 6SKL. **(D)** Close-up of intersubunit PS1-H2 interactions, stall state 1. **(E)** Close-up of intersubunit PS1-H2 interactions, stall state 2.

**Figure S10: Superposition of C-tier states. (A,B)** Superpositions of MCM C-tiers in stall state 1 (grey) and stall state 2 (colored by subunit) (A) and MCM C-tiers in normal fork conformation 1 (PDB 6SKL; grey) and normal fork conformation 2 (PDB 6SKO; colored by subunit) (A), aligned on Mcm5. Spheres correspond to Mcm2 S706. **(C)** Superposition of MCM rings in stall state 1 (grey) and stall state 2 (colored by subunit), aligned on MCM N-tier.

**Figure S11: G4 stall in the reconstituted human DNA replication system. (A)** Purified human replication proteins. **(B)** Superposition of DNA in G4 stall state 1 (green) and human CMG (blue). **(C)** Protein-DNA contacts in human CMG stalled at a leading strand G4. Schematic is analogous to those in Figure S7. **(D)** MCM PS1/H2I loops in human MCM-FPC. Mcm2 is shown in blue, Mcm6, -4, -3, and -7 are shown in green and Mcm5 is shown in orange. **(E)** 5' exonuclease protection analysis to determine position of the yeast CMG-FPC N-tier on DNA substrate harboring a G4 on the leading strand template. The template is the same as in Figure 2B. Green: CMG. Yellow: TIM-TIPIN (TT).

**Figure S12: Local resolution estimations of yeast and human CMG cryo-EM local refinements and consensus maps. (A)** Non-signal-subtracted consensus and local refinements of yeast CMG, G4 stall state 1, colored by local resolution. **(B)** Non-signal-subtracted consensus and local refinements of yeast CMG, G4 stall state 2, colored by local resolution. **(C)** Signal-subtracted consensus and local refinements of human CMG, G4 stall state, colored by local resolution.

### Figure S1

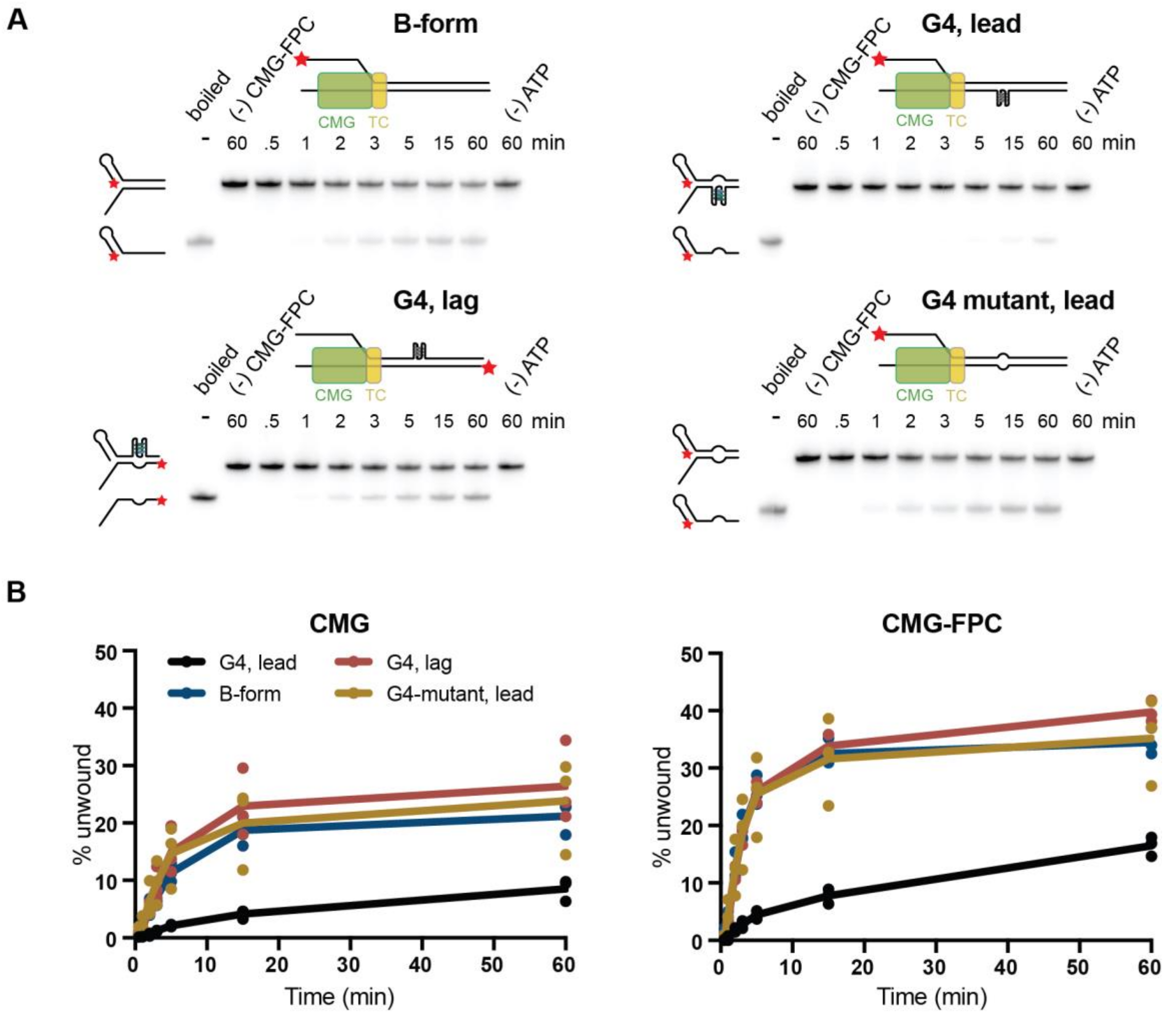
A
B-form
G4, lead
G4, lag
G4 mutant, lead
boiled
(-) CMG-FPC
(-) ATP
CMG
TC
- 60 .5 1 2 3 5 15 60 60 min
B
CMG
CMG-FPC
G4, lead
G4, lag
B-form
G4-mutant, lead
% unwound
Time (min)

### Figure S2

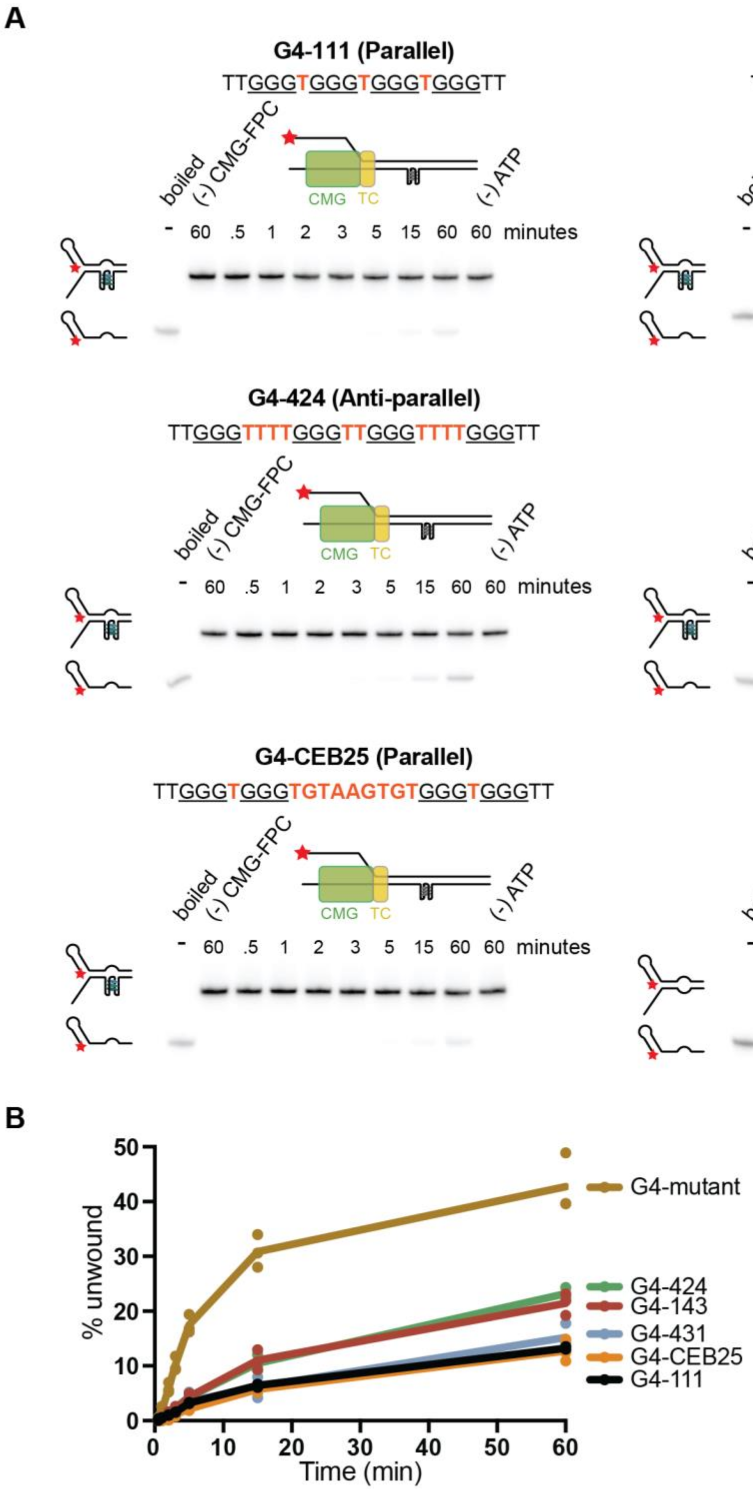
A
G4-111 (Parallel)
TTGGGTGGGTGGGTGGGTT
G4-431 (Parallel)
TTGGGTTTTGGGTTTGGGTGGGTT
G4-424 (Anti-parallel)
TTGGGTTTTGGGTTGGGTTTTGGGTT
G4-143 (Hybrid)
TTGGGTGGGTTTTGGGTTTGGGTT
G4-CEB25 (Parallel)
TTGGGTGGGTGTAAGTGTGGGTGGGTT
G4 mutant
TTGGGTGGGTCCCTGGGTT
boiled
(-) CMG-FPC
CMG
TC
(-) ATP
- 60 .5 1 2 3 5 15 60 60 minutes
B
% unwound
Time (min)
G4-mutant
G4-424
G4-143
G4-431
G4-CEB25
G4-111

### Figure S3

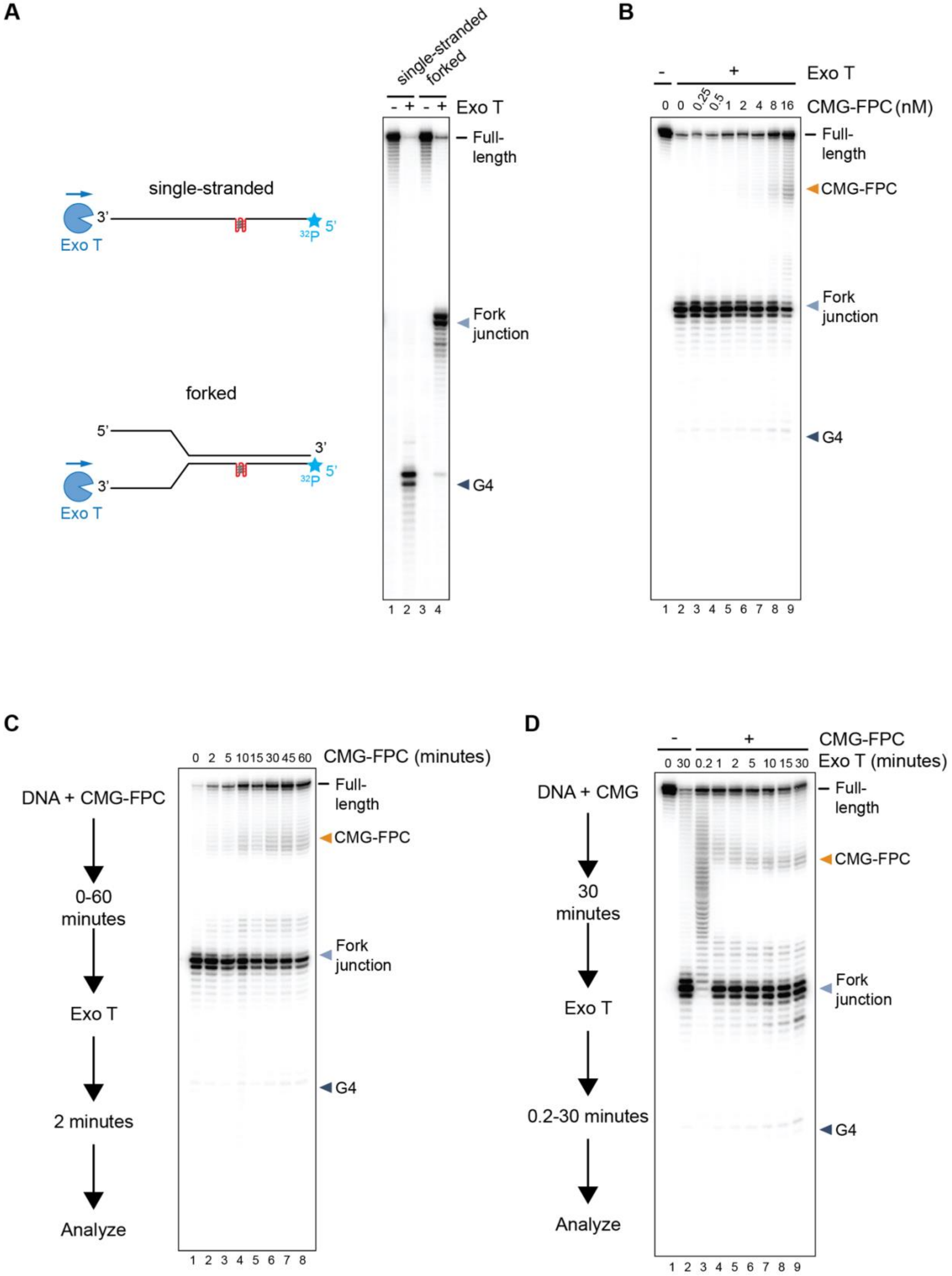
A
single-stranded
3'
5'
Exo T
forked
5'
3'
3'
5'
Exo T
single-stranded
forked
- + - +
Exo T
Full-length
Fork junction
G4
1 2 3 4
B
- +
Exo T
0 0 0.25 0.5 1 2 4 8 16
CMG-FPC (nM)
Full-length
CMG-FPC
Fork junction
G4
1 2 3 4 5 6 7 8 9
C
DNA + CMG-FPC
0-60 minutes
Exo T
2 minutes
Analyze
0 2 5 10 15 30 45 60
CMG-FPC (minutes)
Full-length
CMG-FPC
Fork junction
G4
1 2 3 4 5 6 7 8
D
DNA + CMG
30 minutes
Exo T
0.2-30 minutes
Analyze
- +
CMG-FPC
0 30 0.2 1 2 5 10 15 30
Exo T (minutes)
Full-length
CMG-FPC
Fork junction
G4
1 2 3 4 5 6 7 8 9

### Figure S4

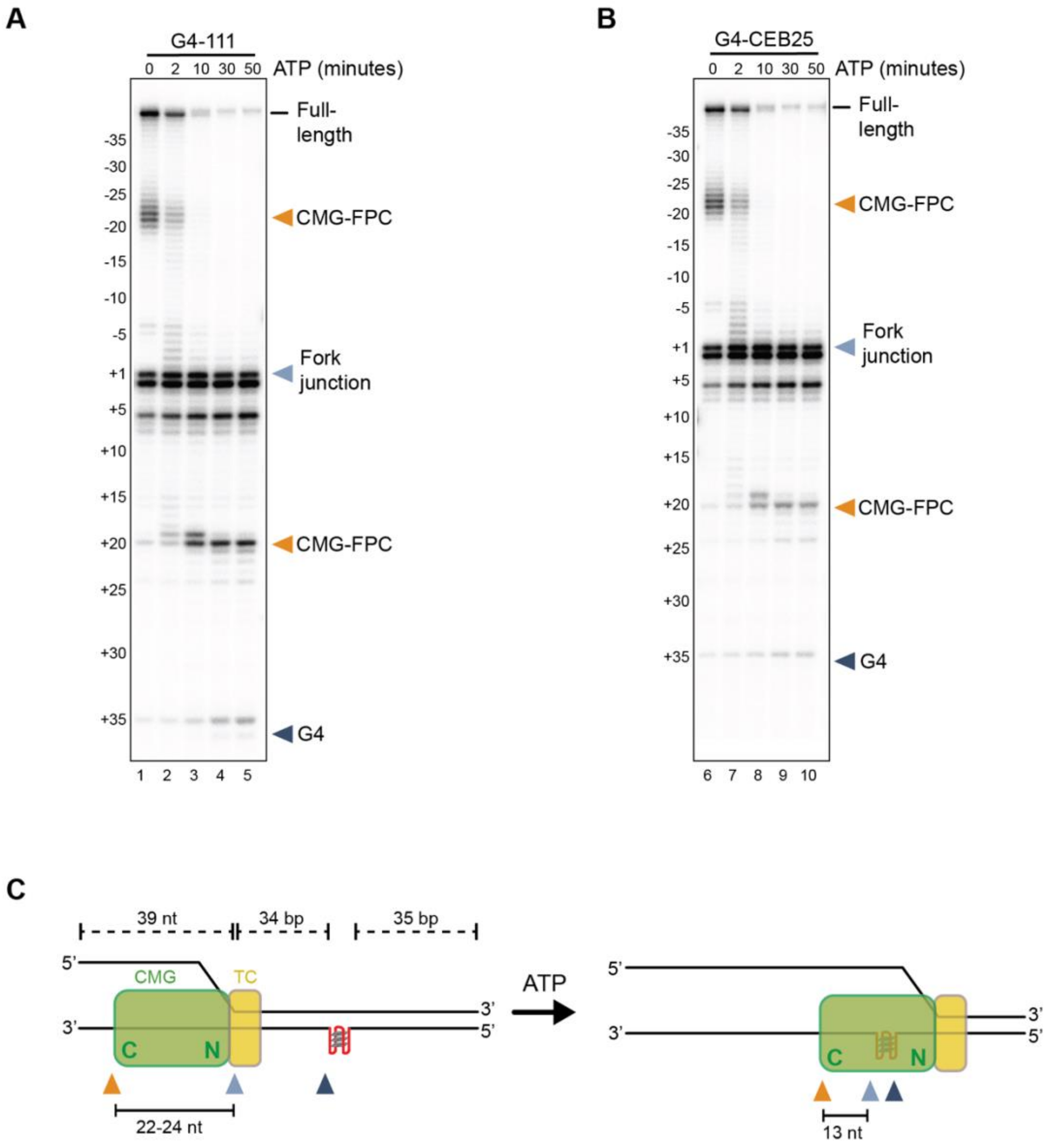
A
G4-111
0 2 10 30 50 ATP (minutes)
Full-length
CMG-FPC
Fork junction
CMG-FPC
G4
-35
-30
-25
-20
-15
-10
-5
+1
+5
+10
+15
+20
+25
+30
+35
1 2 3 4 5
B
G4-CEB25
0 2 10 30 50 ATP (minutes)
Full-length
CMG-FPC
Fork junction
CMG-FPC
G4
6 7 8 9 10
C
39 nt
34 bp
35 bp
5'
3'
CMG
TC
C
N
22-24 nt
ATP
13 nt

### Figure S5

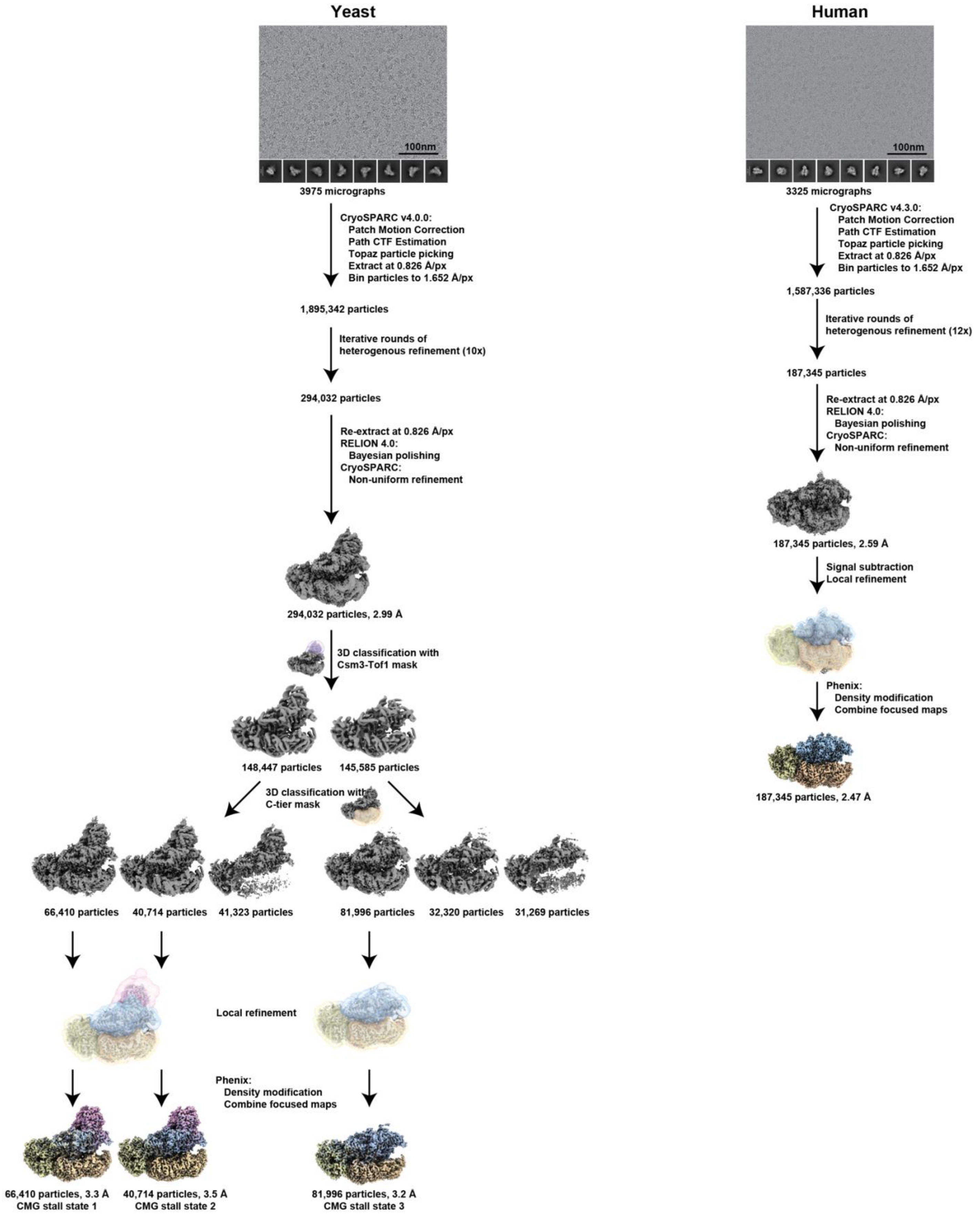
Yeast
100nm
3975 micrographs
CryoSPARC v4.0.0:
Patch Motion Correction
Path CTF Estimation
Topaz particle picking
Extract at 0.826 Å/px
Bin particles to 1.652 Å/px
1,895,342 particles
Iterative rounds of
heterogenous refinement (10x)
294,032 particles
Re-extract at 0.826 Å/px
RELION 4.0:
Bayesian polishing
CryoSPARC:
Non-uniform refinement
294,032 particles, 2.99 Å
3D classification with
Csm3-Tof1 mask
148,447 particles
145,585 particles
3D classification with
C-tier mask
66,410 particles
40,714 particles
41,323 particles
81,996 particles
32,320 particles
31,269 particles
Local refinement
Phenix:
Density modification
Combine focused maps
66,410 particles, 3.3 Å
CMG stall state 1
40,714 particles, 3.5 Å
CMG stall state 2
81,996 particles, 3.2 Å
CMG stall state 3
Human
100nm
3325 micrographs
CryoSPARC v4.3.0:
Patch Motion Correction
Path CTF Estimation
Topaz particle picking
Extract at 0.826 Å/px
Bin particles to 1.652 Å/px
1,587,336 particles
Iterative rounds of
heterogenous refinement (12x)
187,345 particles
Re-extract at 0.826 Å/px
RELION 4.0:
Bayesian polishing
CryoSPARC:
Non-uniform refinement
187,345 particles, 2.59 Å
Signal subtraction
Local refinement
Phenix:
Density modification
Combine focused maps
187,345 particles, 2.47 Å

### Figure S6

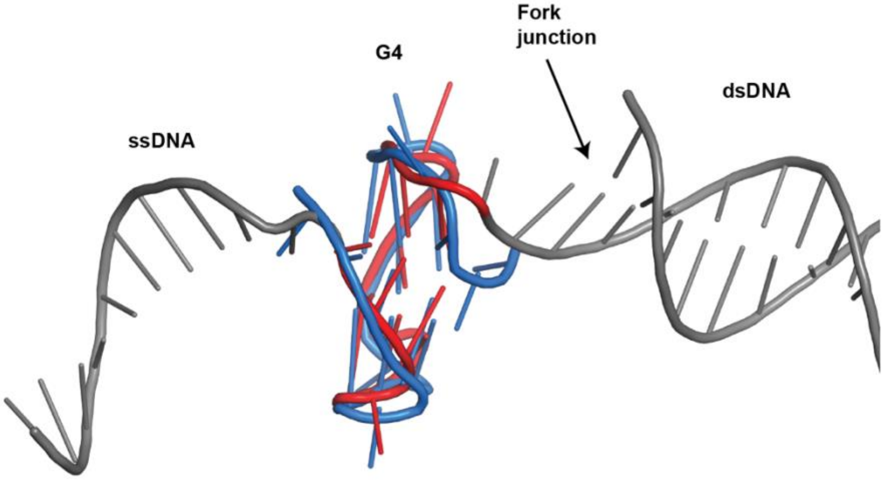

**Figure S7**

State 1

State 2

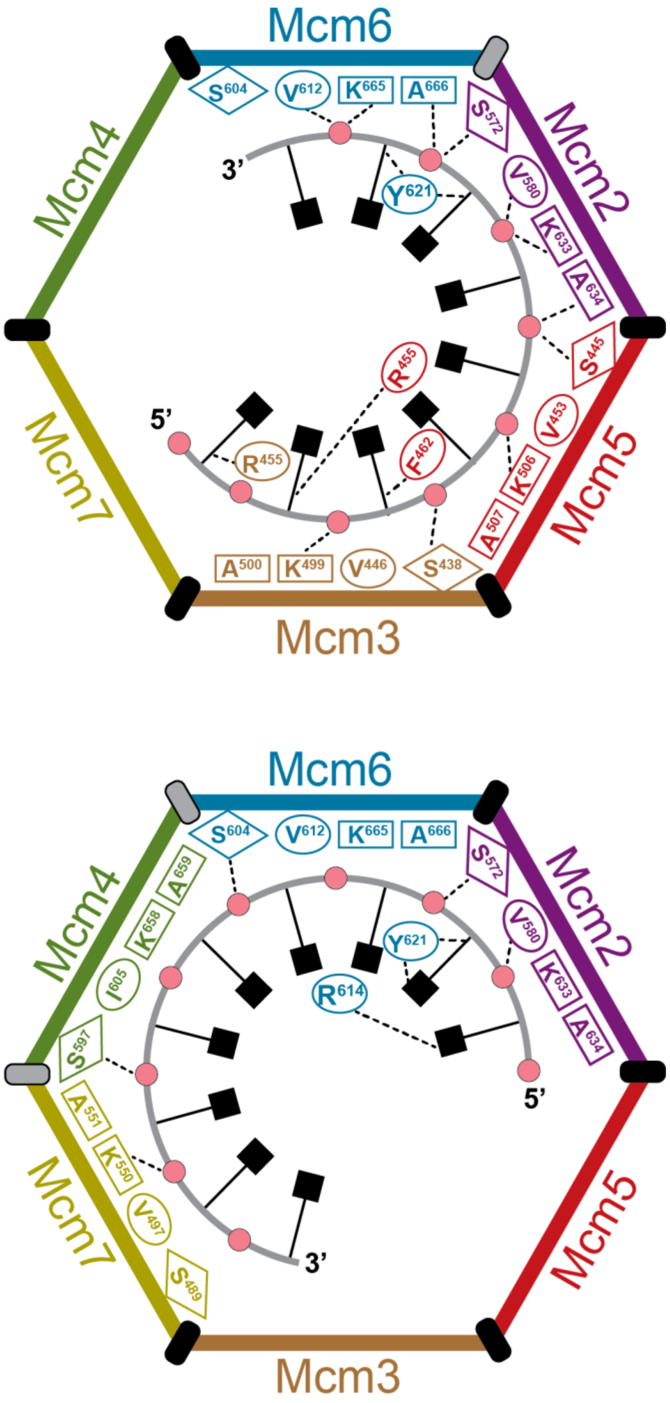

### Figure S8

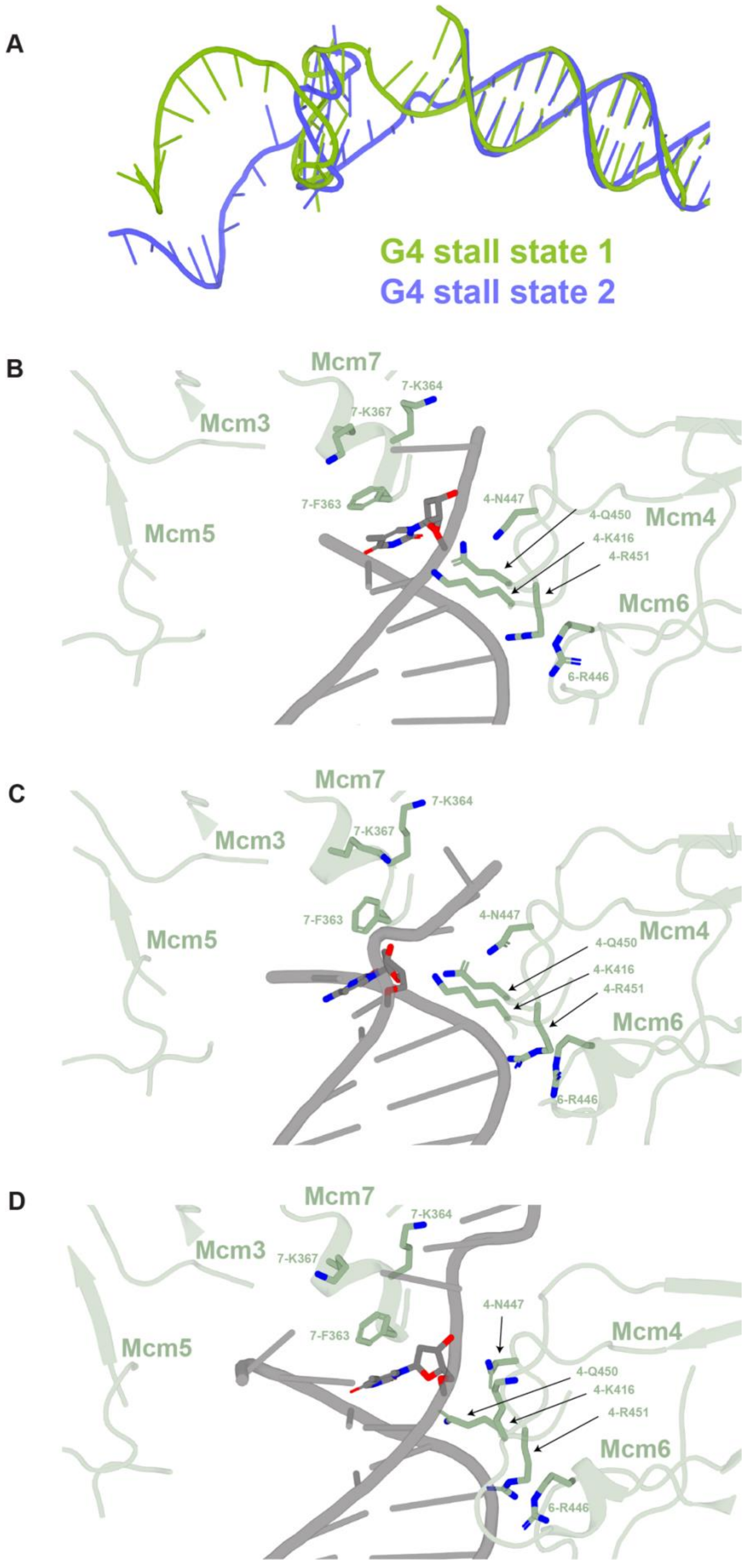
A
G4 stall state 1
G4 stall state 2
B
Mcm7
7-K364
7-K367
Mcm3
Mcm5
7-F363
4-N447
4-Q450
4-K416
4-R451
Mcm4
Mcm6
6-R446
C
Mcm7
7-K364
7-K367
Mcm3
Mcm5
7-F363
4-N447
Mcm4
4-Q450
4-K416
4-R451
Mcm6
6-R446
D
Mcm7
7-K364
Mcm3
7-K367
Mcm5
7-F363
4-N447
Mcm4
4-Q450
4-K416
4-R451
Mcm6
6-R446

### Figure S9

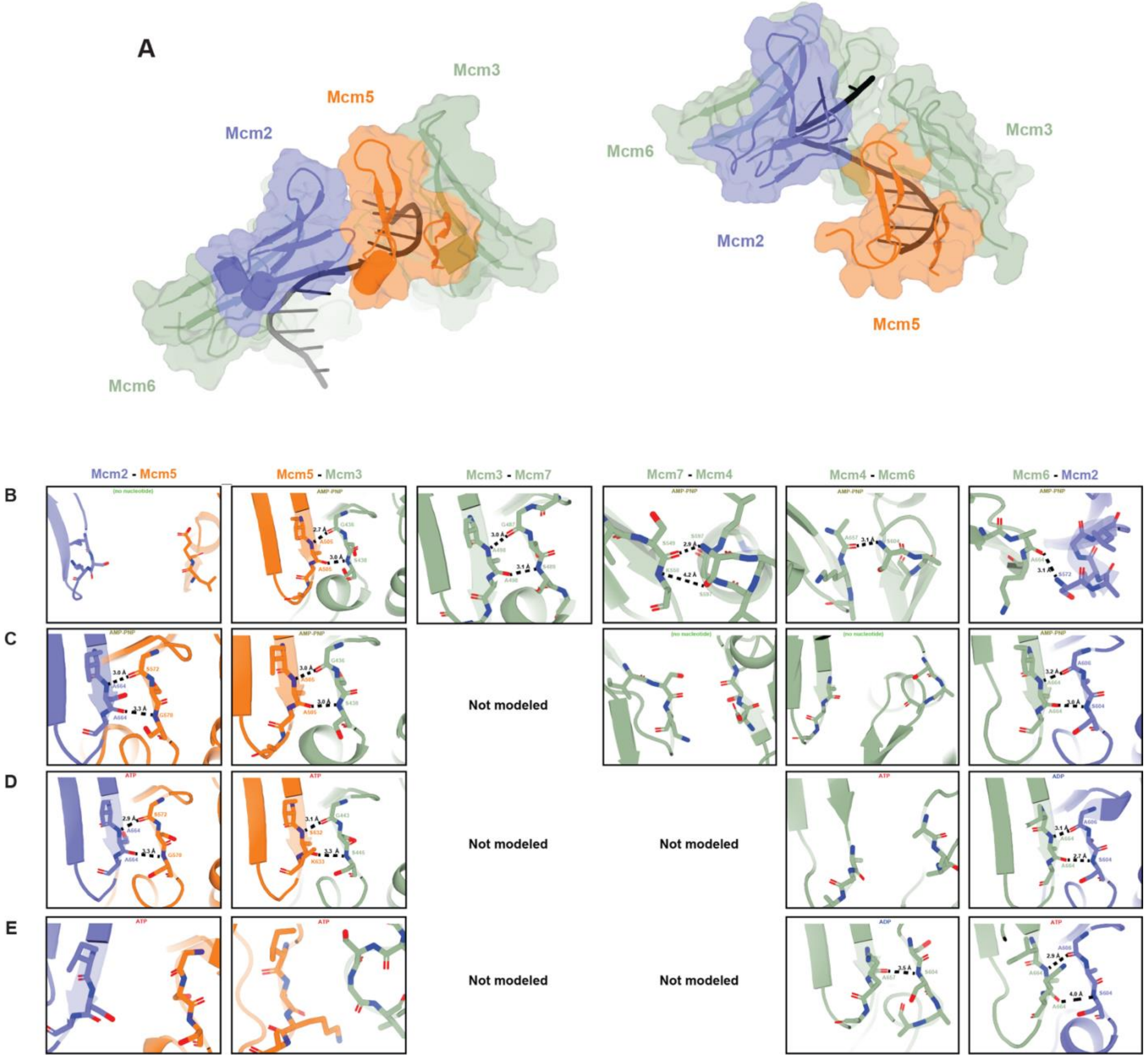

A
Mcm2
Mcm5
Mcm3
Mcm6
Mcm2 - Mcm5
Mcm5 - Mcm3
Mcm3 - Mcm7
Mcm7 - Mcm4
Mcm4 - Mcm6
Mcm6 - Mcm2
B
C
D
E
Not modeled
Not modeled
Not modeled
Not modeled
Not modeled

#### Figure S10

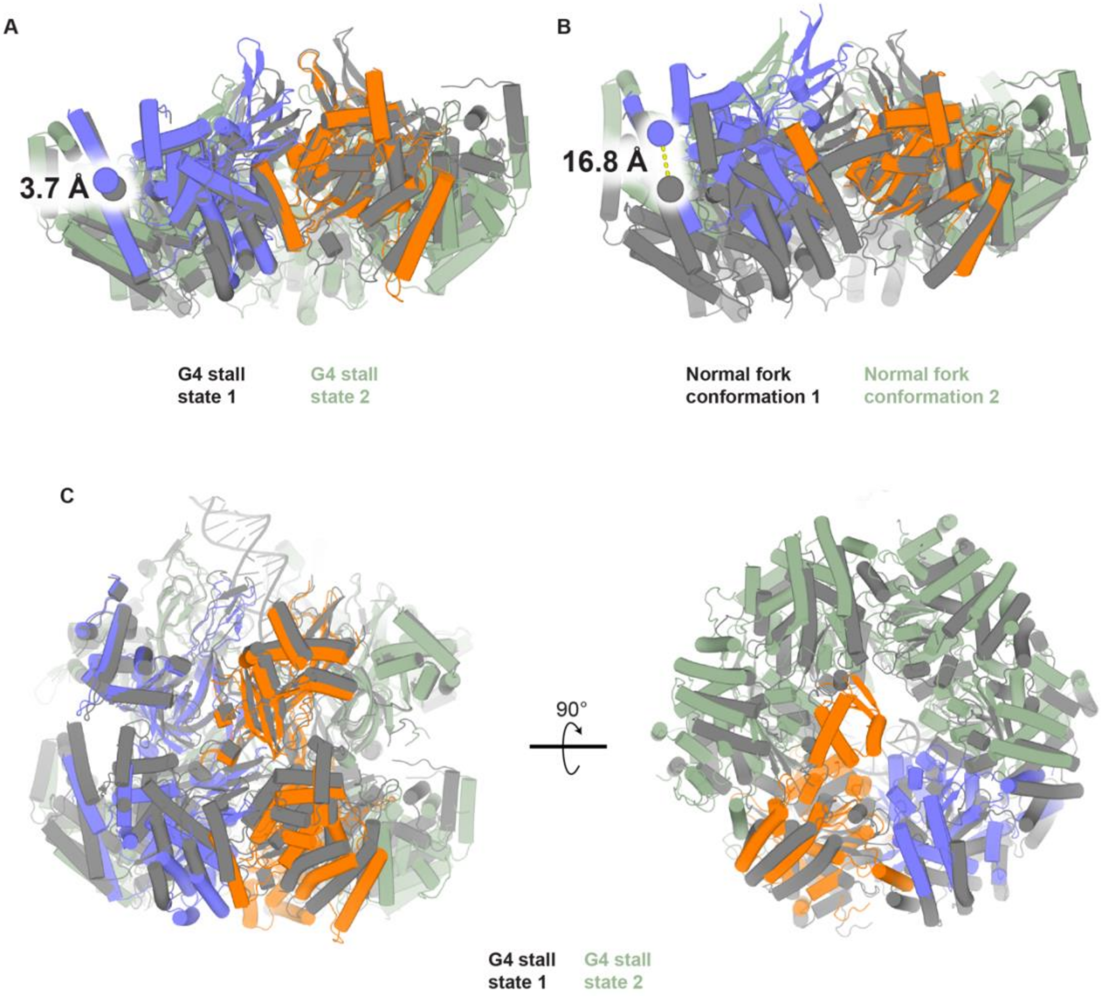
A
3.7 Å
G4 stall state 1
G4 stall state 2
B
16.8 Å
Normal fork conformation 1
Normal fork conformation 2
C
90°
G4 stall state 1
G4 stall state 2

### Figure S11

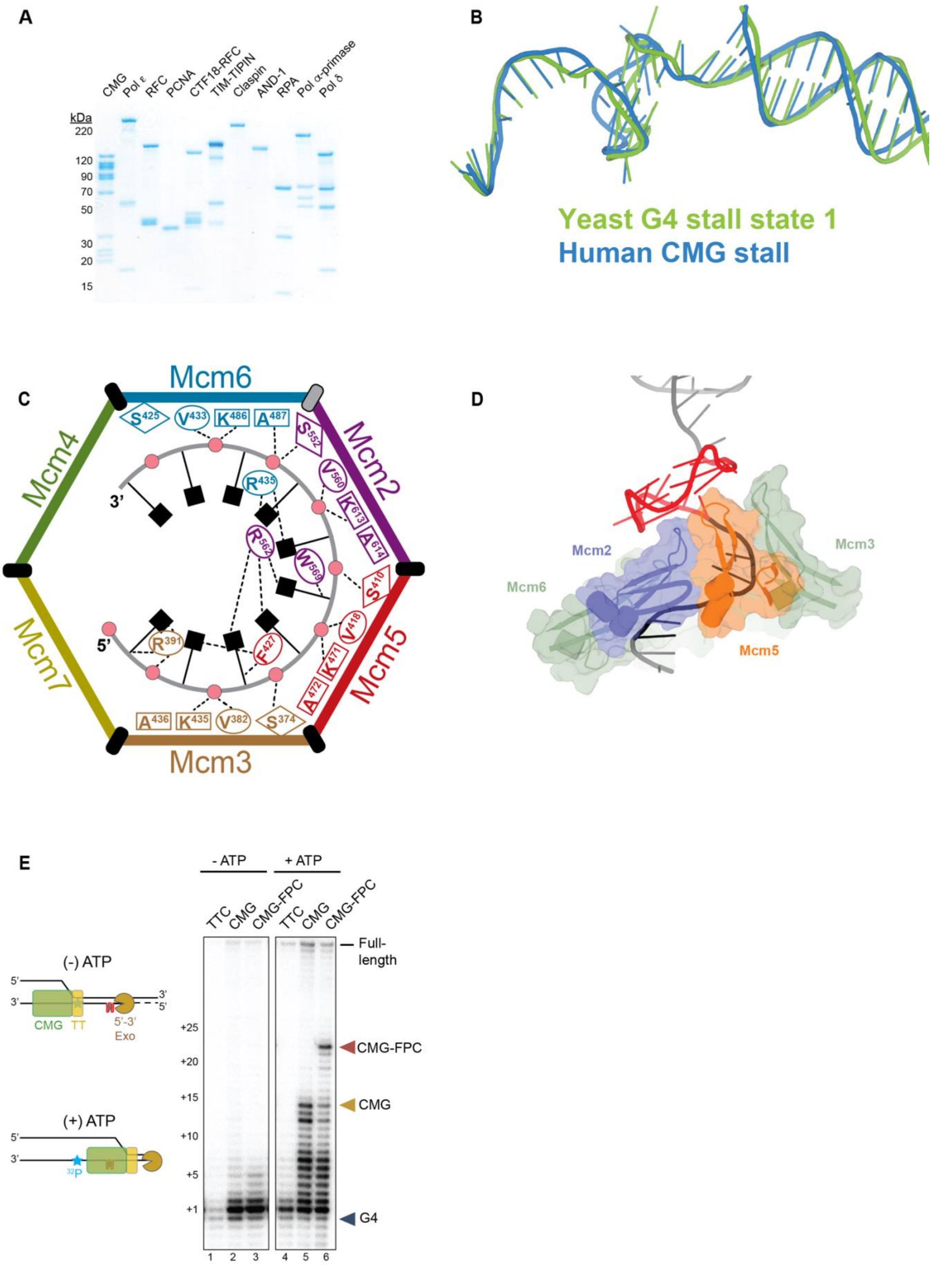
A
CMG
Pol ε
RFC
PCNA
CTF18-RFC
TIM-TIPIN
Claspin
AND-1
RPA
Pol α-primase
Pol δ
kDa
220
120
90
70
50
30
20
15
B
Yeast G4 stall state 1
Human CMG stall
C
Mcm6
Mcm4
Mcm2
Mcm7
Mcm5
Mcm3
3'
5'
D
Mcm2
Mcm3
Mcm6
Mcm5
E
- ATP
+ ATP
TTC
CMG
CMG-FPC
(-) ATP
CMG
TT
5'-3' Exo
(+) ATP
32P
Full-length
CMG-FPC
CMG
G4
+25
+20
+15
+10
+5
+1
1
2
3
4
5
6

### Figure S12

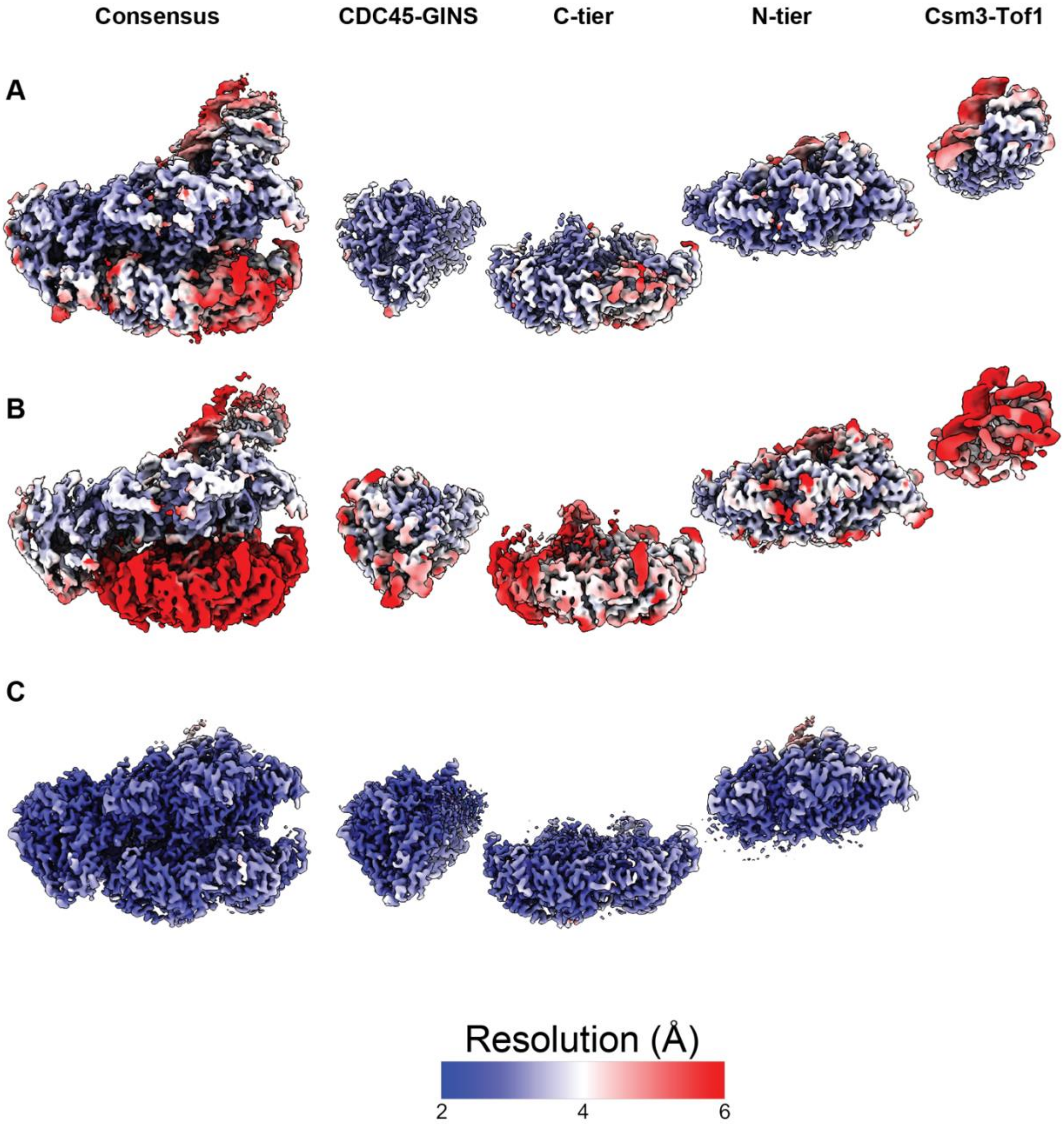
Consensus
CDC45-GINS
C-tier
N-tier
Csm3-Tof1
A
B
C
Resolution (Å)
2
4
6

#### Supplementary Tables

**Table S1: Inserts for yeast/human DNA replication templates.**

| Insert | Oligonucleotides |
|---|---|
| G4, lead | 2984+2985 |
| G4, lag | 3045+3046 |
| B-form | 3043+3044 |

**Table S2: Oligonucleotides for preparing DNA fork of human DNA replication template.**

| Oligonucleotide | Purpose |
|---|---|
| 3097 | Tracking strand with phosphorylated 5' end for ligation. |
| 3128 | Non-tracking strand. |
| 2529 | Primer for leading strand synthesis. |

**Table S3: Oligonucleotides to prepare substrate for helicase and 3' exonuclease protection assays.**

| Template | Tracking strand | Non-tracking strand | Figure |
|---|---|---|---|
| G4, lead | 2883 | 2884 | Figure 2A, Suppl. Figure 1, Suppl. Figure 3A. |
| G4 mutant, lead | 3173 | 2884 | Suppl. Figure 1. |
| G4, lag | 3174 | 3175 | Suppl. Figure 1. |
| B-form | 2883 | 2894 | Suppl. Figure 1. |
| G4-111 | 3098 | 3099 | Suppl. Figure 2, Suppl. Figure 4. |
| G4-431 | 3158 | 3159 | Suppl. Figure 2. |
| G4-424 | 3156 | 3157 | Suppl. Figure 2. |
| G4-143 | 3154 | 3155 | Suppl. Figure 2. |
| G4-CEB25 | 3152 | 3153 | Suppl. Figure 2, Suppl. Figure 4. |
| G4 mutant | 3177 | 3099 | Suppl. Figure 2. |

| | | | |
|---|---|---|---|
| G4, lead | 2560 | 2559 | Suppl. Figure 3B-D. |

**Table S4: List of Oligonucleotides**

| **Name** | **Sequence** |
|---|---|
| 2529 | CCTCTCGAGCCCATCCTTCCACTTCCCAACCCTCACC |
| 2559 | GGCAGGCAGGCAGGCAGGCAGGCAGGCAGGCAGGCAGGCAGGTTGGCGAAT TCCCTTTTTTTTTTTTTTTTTTTTCCTCAGCACGACGTTGTAAAACGAG |
| 2560 | GGCTCGTTTTACAACGTCGTGCTGAGGTTGGGTGGGTGGGTGGGTTGGGAAT TCGCCAACCTTTTTTTTTTTTTTTTTTTTTTTTTTTTTTTTTTTTTTTTT |
| 2602 | TTTTTTTTTTTTTTTTTTTTTTTTTTTTTTTTTTTTTTTT |
| 2883 | CTCGTTTTACAACGTCGTGCTGAGTGATATCTGCTTTGGGTGGGTGGGTGGGT TGAGGCAATCTGAATTCGCCAACCTTTTTTTTTTTTTTTTTTTTTTTTTTTTTTTT TTTTTTTT |
| 2884 | GGCAGGCAGGCAGGCAGGCAGGCAGGCAGGCAGGCAGGCAGGTTGGCGAAT TCAGATTGCCTCTTTTTTTTTTTTTTTTTTAGCAGATATCACTCAGCACGACGT TGTAAAACGAG |
| 2894 | GGCAGGCAGGCAGGCAGGCAGGCAGGCAGGCAGGCAGGCAGGTTGGCGAAT TCAGATTGCCTCAACCCACCCACCCACCCAAAGCAGATATCACTCAGCACGAC GTTGTAAAACGAG |
| 2970 | ACCTTTTTTTTTTTTTTTTTTTTTTTTTTTTTTTTTTTTTTTT |
| 2973 | rArArArArArArArArArArGrGrUrUrGrGrCrGrArArUrUrCrArG |
| 2984 | /5Phos/GAATCTCGTTTTACACGTGCGTGCTGAGTGATATCTGCTTTGGGTGGGT GGGTGGGTTGAGGCAATCTGAAGTATACTTCGCCAACCT |
| 2985 | /5Phos/TGAGAGGTTGGCGAAGTATACTTCAGATTGCCTCTTTTTTTTTTTTTTT TTTTAGCAGATATCACTCAGCACGCACGTGTAAAACGAG |
| 2990 | GGCTCGTTTTACAACGTCGTGCTGAGTGATATCTGCTTTGGGTGGGTGGGTGG GTTGAGGCAATCTGAATTCGCCA |
| 3043 | /5Phos/GAATCTCGTTTTACACGTGCGTGCTGAGTGATATCTGCTTTACCGTAGG CTGTGACTTGAGGCAATCTGAAGTATACTTCGCCAACCT |
| 3044 | /5Phos/TGAGAGGTTGGCGAAGTATACTTCAGATTGCCTCAAGTCACAGCCTAC GGTAAAGCAGATATCACTCAGCACGCACGTGTAAAACGAG |
| 3045 | /5Phos/TGAGCTCGTTTTACACGTGCGTGCTGAGTGATATCTGCTTTGGGTGGGT GGGTGGGTTGAGGCAATCTGAAGTATACTTCGCCAACCT |
| 3046 | /5Phos/GAATAGGTTGGCGAAGTATACTTCAGATTGCCTCTTTTTTTTTTTTTTT TTTAGCAGATATCACTCAGCACGCACGTGTAAAACGAG |

| | |
|---|---|
| 3097 | /5Phos/GGCCGAGGAAGCCATGGTGAGAATTGGAGAGTGTGTTTTTTTTTTTTT TTTTTTTTTTTTTTTTTTTTTTTTTGAGGAAAGAATGTTGGTGAGGGTTGGGA AGTGGAAGGATGGGCTCGAGAGGTTTTTTTTTTTTTTTTTTTTTTTTTTT*T *T*T*T |
| 3098 | CTCGTTTTACAACGTCGTGCTGAGTGATATCTGCTTTGGGTGGGTGGGTGGGT TCTGTTCGCTCGAGGCAATCTGAATTCGCCAACCTTTTTTTTTTTTTTTTTTT TTTTTTTTTTTTTTTTTT |
| 3099 | GGCAGGCAGGCAGGCAGGCAGGCAGGCAGGCAGGCAGGCAGGTTGGCGAAT TCAGATTGCCTCGAGCGAACAGTTTTTTTTTTTTTTTTTTAGCAGATATCACTC AGCACGACGTTGTAAAACGAG |
| 3128 | GGCAGGCAGGCAGGCAGGCAGGCAGGCAGGCAGGCAGGCACACACTCTCCA ATTCTCACCATGGCTTCCTCGGCCCTGA |
| 3152 | CTCGTTTTACAACGTCGTGCTGAGTGATATCTGCTTTGGGTGGGTGTAAGTGT GGGTGGGTTCTGTTCGCTCGAGGCAATCTGAATTCGCCAACCTTTTTTTTTTT TTTTTTTTTTTTTTTTTTTTTTTTTTT |
| 3153 | GGCAGGCAGGCAGGCAGGCAGGCAGGCAGGCAGGCAGGCAGGTTGGCGAAT TCAGATTGCCTCGAGCGAACAGTTTTTTTTTTTTTTTTTTTTTTTTTTTAGCAGA TATCACTCAGCACGACGTTGTAAAACGAG |
| 3154 | CTCGTTTTACAACGTCGTGCTGAGTGATATCTGCTTTGGGTGGGTTTTGGGTTT GGGTTCTGTTCGCTCGAGGCAATCTGAATTCGCCAACCTTTTTTTTTTTTTTT TTTTTTTTTTTTTTTTTTTTTTTT |
| 3155 | GGCAGGCAGGCAGGCAGGCAGGCAGGCAGGCAGGCAGGCAGGTTGGCGAAT TCAGATTGCCTCGAGCGAACAGTTTTTTTTTTTTTTTTTTTTTTTTTTAGCAGATAT CACTCAGCACGACGTTGTAAAACGAG |
| 3156 | CTCGTTTTACAACGTCGTGCTGAGTGATATCTGCTTTGGGTTTTGGGTTGGGTT TTGGGTTCTGTTCGCTCGAGGCAATCTGAATTCGCCAACCTTTTTTTTTTTTT TTTTTTTTTTTTTTTTTTTTTTTTTT |
| 3157 | GGCAGGCAGGCAGGCAGGCAGGCAGGCAGGCAGGCAGGCAGGTTGGCGAAT TCAGATTGCCTCGAGCGAACAGTTTTTTTTTTTTTTTTTTTTTTTTTTTAGCAGAT ATCACTCAGCACGACGTTGTAAAACGAG |
| 3158 | CTCGTTTTACAACGTCGTGCTGAGTGATATCTGCTTTGGGTTTTGGGTTTGGGT GGGTTCTGTTCGCTCGAGGCAATCTGAATTCGCCAACCTTTTTTTTTTTTTTT TTTTTTTTTTTTTTTTTTTTTTTT |
| 3159 | GGCAGGCAGGCAGGCAGGCAGGCAGGCAGGCAGGCAGGCAGGTTGGCGAAT TCAGATTGCCTCGAGCGAACAGTTTTTTTTTTTTTTTTTTTTTTTTTTAGCAGATAT CACTCAGCACGACGTTGTAAAACGAG |

| | |
|---|---|
| 3173 | CTCGTTTTACAACGTCGTGCTGAGTGATATCTGCTTTGGGTGGGTCCCTGGGTTGAGGCAATCTGAATTCGCCAACCTTTTTTTTTTTTTTTTTTTTTTTTTTTTTTTTTTTTT |
| 3174 | CTCGTTTTACAACGTCGTGCTGAGTGATATCTGCTTTTTTTTTTTTTTTTTTTGAGGCAATCTGAATTCGCCAACCTTTTTTTTTTTTTTTTTTTTTTTTTTTTTTTTTTTTT |
| 3175 | GGCAGGCAGGCAGGCAGGCAGGCAGGCAGGCAGGCAGGCAGGTTGGCGAATTCAGATTGCCTCTTGGGTGGGTGGGTGGGTTAGCAGATATCACTCAGCACGACGTTGTAAAACGAG |
| 3177 | CTCGTTTTACAACGTCGTGCTGAGTGATATCTGCTTTGGGTGGGTCCCTGGGTTCTGTTCGCTCGAGGCAATCTGAATTCGCCAACCTTTTTTTTTTTTTTTTTTTTTTTTTTTTTTTTTTTTT |

/5Phos/: 5' phosphate, r: ribose, *: phosphorothioate

**Table S5: List of yeast strains**

| **Strain** | **Reference** |
|---|---|
| yJF38 (Cdt1-Mcm2-7 purification) | (*81*) |
| ySDORC (ORC purification) | (*81*) |
| ySA35 (DDK purification) | (*65*) |
| ySD13 (Sld2 purification) | (*66*) |
| ySD15 (Cdc45 purification) | (*66*) |
| yDR110 (Dpb11 purification | (*66*) |
| yDR116 (Pol ε purification) | (*66*) |
| yDR105 (CDK purification) | (*66*) |
| ySD16 (Pol α purification) | (*66*) |
| yDR163 (Mrc1 purification) | (*66*) |
| yDR137 (Csm3-Tof1 purification) | (*66*) |
| yIW389 (RFC purification) | (*66*) |
| yDR131 (Pol δ purification) | (*66*) |
| yDR109 (GINS purification) | (*66*) |
| yDR128 (Top1 purification) | (*66*) |
| ySD35 (CMG purification) | (*44*) |
| yCL37 (hRFC purification) Genotype: *MATa ade2-1 ura3-1 his3-11,15 trp1-1 leu2-3,112 can1-* | This study |

| | |
|---|---|
| *100 pep4::kanMX bar::hphNAT1 (hygromycinB) trp1:: 3xFLAG-TEV-GAL-RFC1 (TRP1) leu2::GAL-RFC2/RFC3 (LEU2) ura3::GAL-RFC4/RFC5 (URA3)* | |
| yCL71 (CTF18-RFC purification) Genotype: *MATa ade2-1 ura3-1 his3-11,15 trp1-1 leu2-3,112 can1-100 pep4::kanMX bar::hphNAT1 (hygromycinB) leu2::GAL-RFC2/RFC3 (LEU2) ura3:: RFC4/RFC5 (URA3) trp1:: 3xFLAG-TEV-CHTF18 (TRP1) his3:: DSCC1-CHTF8 (HIS3)* | This study |

**Table S6: List of plasmids**

| **Plasmid** | **Reference** |
|---|---|
| p75 (Bacterial expression of Cdc6) | (*82*) |
| p1280 (Bacterial expression of Sld3) | This study |
| p1281 (Bacterial expression of Sld7) | This study |
| p801 (Bacterial expression of Ctf4) | (*66*) |
| pJM126 (Bacterial expression of RPA) | (*83*) |
| p1274 (Bacterial expression of Mcm10) | This study |
| p1012 (Bacterial expression of PCNA) | (*66*) |
| p1345 (Baculovirus generation for human CDC45 expression) | This study |
| p1378 (Baculovirus generation for CLASPIN expression) | This study |
| p1379 (Baculovirus generation for AND1 expression) | This study |
| p1356 (Baculovirus generation for human GINS expression) | This study |
| p1358 (Baculovirus generation for human MCM2-7 expression) | This study |
| p1396 (Baculovirus generation for human Pol e expression) | This study |
| p1404 (Baculovirus generation for TIMELESS-TIPIN expression) | This study |

| | |
|---|---|
| p1463 (Baculovirus generation for human Pol d expression) | This study |
| p1477 (Baculovirus generation for human Pol a expression) | This study |
| p1361 (Bacterial expression of human PCNA) | This study |
| p1411 (Bacterial expression of human RPA) | (*68*) |
| p1400 (Template for yeast replication assays) | This study |
| p1432 (Template for human replication assays) | This study |

**Table S7: EM validation and refinement statistics**

| Name | G4 stall state 1 | G4 stall state 2 | G4 stall state 3 | hCMG stall |
|---|---|---|---|---|
| Symmetry imposed | C1 | C1 | C1 | C1 |
| Map resolution (Å) at FSC=0.143 | | | | |
| Consensus | 3.15 | 3.52 | 3.15 | 2.59 |
| MCM N-tier | 3.06 | 3.28 | 3.1 | 2.53 |
| MCM C-tier | 3.15 | 3.61 | 3.1 | 2.46 |
| Cdc45-GINS | 3.12 | 3.45 | 3.11 | 2.41 |
| Tof1-Csm3 | 3.16 | 4.3 | --- | --- |
| Map resolution range (Å) | 3.1-3.2 | 3.3-4.3 | 3.1-3.1 | 2.4-2.6 |
| Model resolution (Å) at FSC=0.5 | 3.1 | 3.4 | 3.1 | 2.2 |
| Map-model CC (masked) | 0.83 | 0.81 | 0.84 | 0.87 |
| Model composition | 48564 | 49288 | 41869 | 40996 |
| Non-hydrogen atoms | 5883 | 5977 | 5129 | 4975 |
| Protein residues | 69 | 68 | 42 | 53 |
| Nucleotides | 17 | 16 | 17 | 18 |
| Ligands | 48564 | 49288 | 41869 | 40996 |
| Mean B-factors (Å$^2$) | 88.58 | 81.75 | 87.76 | 50 |
| Protein | 123.72 | 106.67 | 97.8 | 49.95 |
| Nucleotides | 98.64 | 81.88 | 87.35 | 40.07 |

| | | | | |
|---|---|---|---|---|
| Ligands | 88.58 | 81.75 | 87.76 | 50 |
| Validation | | | | |
| MolProbability Score | 1.37 | 1.89 | 1.68 | 1.13 |
| Clashscore | 4.22 | 7.07 | 5.46 | 3.42 |
| Poor rotamers (%) | 1.33 | 2.24 | 2.17 | 0.89 |
| | 1.37 | 1.89 | 1.68 | 1.13 |
| Ramachandran Plot | | | | |
| Disallowed (%) | 0 | 0.03 | 0 | 0.02 |
| Allowed (%) | 2.31 | 3.55 | 2.66 | 1.88 |
| Favored (%) | 97.69 | 96.41 | 97.34 | 98.1 |
